# A conserved choreography of mRNAs at centrosomes reveals a localization mechanism involving active polysome transport

**DOI:** 10.1101/2020.09.04.282038

**Authors:** Adham Safieddine, Emeline Coleno, Abdel-Meneem Traboulsi, Oh Sung Kwon, Frederic Lionneton, Virginie Georget, Marie-Cécile Robert, Thierry Gostan, Charles Lecellier, Soha Salloum, Racha Chouaib, Xavier Pichon, Hervé Le Hir, Kazem Zibara, Marion Peter, Edouard Bertrand

## Abstract

Local translation allows for a spatial control of gene expression. Here, we used high-throughput smFISH to screen centrosomal protein-coding genes, and we describe 8 human mRNAs accumulating at centrosomes. These mRNAs localize at different stages during cell cycle with a remarkable choreography, indicating a finely regulated translational program at centrosomes. Interestingly, drug treatments and reporter analyses revealed a common translation-dependent localization mechanism requiring the nascent protein. Using ASPM and NUMA1 as models, single mRNA and polysome imaging revealed active movements of endogenous polysomes towards the centrosome at the onset of mitosis, when these mRNAs start localizing. ASPM polysomes associate with microtubules and localize by either motor-driven transport or microtubule pulling. Remarkably, the *Drosophila* orthologs of the human centrosomal mRNAs also localize to centrosomes and also require translation. These data identify a conserved family of centrosomal mRNAs that localize by active polysomes transport mediated by nascent proteins.

## Introduction

Messenger RNA localization is a post-transcriptional process by which cells target certain mRNAs to specific subcellular compartments. The trafficking of mRNA molecules is linked to its metabolism and function (Kejiou and Palazzo, 2017). Indeed, the subcellular localization of a transcript can influence its maturation, translation, and degradation. On one hand, mRNAs can be stored in a translationally repressed state in dedicated structures such as P-bodies (Hubstenberger et al., 2017). On the other hand, some mRNAs can localize to be translated locally. Such a local protein synthesis can be used to localize the mature polypeptide, and in this case it can contribute to a wide range of functions such as cell migration, cell polarity, synaptic plasticity, asymmetric cell divisions, embryonic patterning and others (Cody et al., 2013; Chin and Lécuyer, 2017; Ryder and Lerit, 2018 for reviews). Recently, local translation has also been linked to the metabolism of the nascent protein, rather than to localize the mature polypeptide. This is for instance the case for mRNAs translated in distinct foci termed translation factories, which correspond to small cytoplasmic aggregates containing multiple mRNA molecules of a given gene (Pichon et al., 2016; Chouaib et al., 2019).

Specific sub-cellular localization of mRNA molecules can be achieved by several mechanisms. Passive diffusion coupled with local entrapment and/or selective local protection from degradation are two strategies that can establish specific distributions of mRNA molecules (reviewed in Chin and Lécuyer, 2017). In most cases however, mRNA transport and localization occurs via motor-driven transport on the cytoskeleton (Bertrand et al., 1998; Bullock, 2007; Buxbaum et al., 2015). Molecular elements that regulate and control mRNA localization include *cis*- and *trans*-acting elements. *Cis*-acting elements are referred to as zip-codes and are often found within the 3’UTR of the transcript (Trcek and Singer, 2010; Kannaiah and Amster-Choder, 2014; Kim et al., 2015 for reviews). Many types of zip-codes have been described based on primary sequence, number, redundancy, and secondary/tertiary structure. Zip-codes are defined by their ability to carry sufficient information for localizing the transcript. They bind one or several trans-acting RNA-binding proteins (RBPs), which mediate diverse aspects of RNA metabolism such as motor binding and translational regulation (reviewed in Chin and Lécuyer, 2017). Indeed, mRNAs in transit are often subjected to a spatial control of translation (reviewed in Besse and Ephrussi, 2008). A long-standing notion in the field is that the transport of localized mRNAs occurs in a translationally repressed state, which serves to spatially restrict protein synthesis (Vazquez-Pianzola and Suter, 2012; Vazquez-Pianzola et al., 2016). Local translational de-repression occurs once the transcript has reached its destination, for instance by phosphorylation events and/or competition with pre-existing local proteins (Hüttelmaier et al., 2005; Paquin et al., 2007; Götze et al., 2017)

While active transport of transcripts through RNA zip-codes appears to be a frequent mechanism, mRNA localization can also involve the nascent polypeptide, as in the case of secreted proteins. Here, the signal recognition particle (SRP) binds the nascent signal peptide, inhibits translation elongation, and mediates anchoring of the nascent polysome to the SRP receptor on the endoplasmic reticulum, where translation elongation resumes (Gilmore et al., 1982; Walter and Blobel, 1980, 1982). Recently, a few hints, such as puromycin sensitivity, suggested that translation may play a role in the localization of some other types of mRNAs (Sepulveda et al., 2018; Chouaib et al, 2019). Whether this is indeed the case and the mechanisms involved remain however unknown.

Centrosomes are ancient and evolutionary conserved organelles that function as microtubule (MT) organizing centers in most animal cells. They play key roles in cell division, signaling, polarity and motility (Wu and Akhmanova, 2017; Breslow and Holland, 2019; for reviews). A centrosome is composed of two centrioles and their surrounding pericentriolar material (PCM). In cycling cells, centriole duplication is tightly coupled to the cell cycle to ensure a constant number of centrioles in each cell after mitosis (reviewed in Nigg and Holland, 2018). Briefly, G1 cells contain one centrosome with two centrioles connected by a linker. At the beginning of S phase, each parental centriole orthogonally assembles one new procentriole. This configuration is termed engagement and prevents reduplication of the parental centrioles. Procentrioles elongate as the cell is progressing through S and G2. In G2, the two centriolar pairs mature and PCM expands, in preparation of mitotic spindle formation (Breslow and Holland, 2019). The G2/M transition marks the disruption of the centriole linker and centrosome separation. The first clues suggesting the importance of mRNA localization and local translation at the centrosomes were discovered almost 20 years ago in *Xenopus* early embryos (Groisman et al., 2000). It was found that cyclin B mRNAs concentrated on the mitotic spindle, and that this localization was dependent on the ability of CPEB to associate with microtubules and centrosomes. A more global view was obtained in *Drosophila* embryos where a systematic analysis of RNA localization was performed (Lécuyer et al., 2007). Although this study did not reach single molecule sensitivity, it revealed that 6 mRNAs localized at centrosomes across different stages of early *Drosophila* development. In a following study, 13 mRNAs were annotated as enriched on *Drosophila* centrosomes in at least one stage/tissue over the full course of embryogenesis (Wilk et al., 2016). In humans, four mRNAs were recently found to localize at centrosomes (PCNT, ASPM, NUMA1 and HMMR; Sepulveda et al., 2018; Chouaib et al., 2019). All these mRNAs all code for centrosomal proteins, suggesting that they are translated locally.

Here, we performed a systematic single molecule Fluorescent *in situ* hybridization (smFISH) screen of almost all human mRNAs coding for centrosomal proteins and we described a total of 8 transcripts localizing at centrosomes. Remarkably, all 8 mRNAs required synthesis of the nascent protein to localize and, by imaging single ASPM and NUMA1 mRNAs and polysomes, we demonstrate that localization occurs by active transport of polysomes. Moreover, the *Drosophila* orthologs of the human centrosomal mRNAs also localized to centrosomes in a similar translation-dependent manner. This work thus identifies a conserved family of centrosomal mRNAs that become localized by active polysome transport.

## Results

### Screening genes encoding centrosomal proteins reveals a total of 8 human mRNAs localizing at the centrosome

In order to acquire a global view of centrosomal mRNA localization in human cells, we developed a high-throughput smFISH technique (HT-smFISH) and screened genes encoding centrosomal and mitotic spindle proteins. The experimental pipeline is described in Figure 1A. Briefly, we designed 50 to 100 individual probes against each mRNA of the screen. The probes were then generated from complex pools of oligonucleotides (92,000), first by using gene-specific primers to PCR out the probes of single genes, followed by a second round of PCR to add a T7 promoter and *in vitro* transcription to generate single-stranded RNA probes (Figure 1A; see Materials and Methods). The probes were designed such that each contained a gene-specific sequence flanked by two overhangs common to all probes (flaps X and Y). A pre-hybridization step then labeled the overhangs with fluorescently labeled locked nucleic acid (LNA) oligonucleotides, and the heteroduplexes were hybridized on cells as in the smiFISH technique (Tsanov et al., 2016), except that cells were grown and hybridized on 96-well plates. This approach is cost-effective because the probes are generated from an oligonucleotide pool. In addition, the probes can be used individually or combined in different colors, allowing a flexible experimental design.

**Figure 1:**
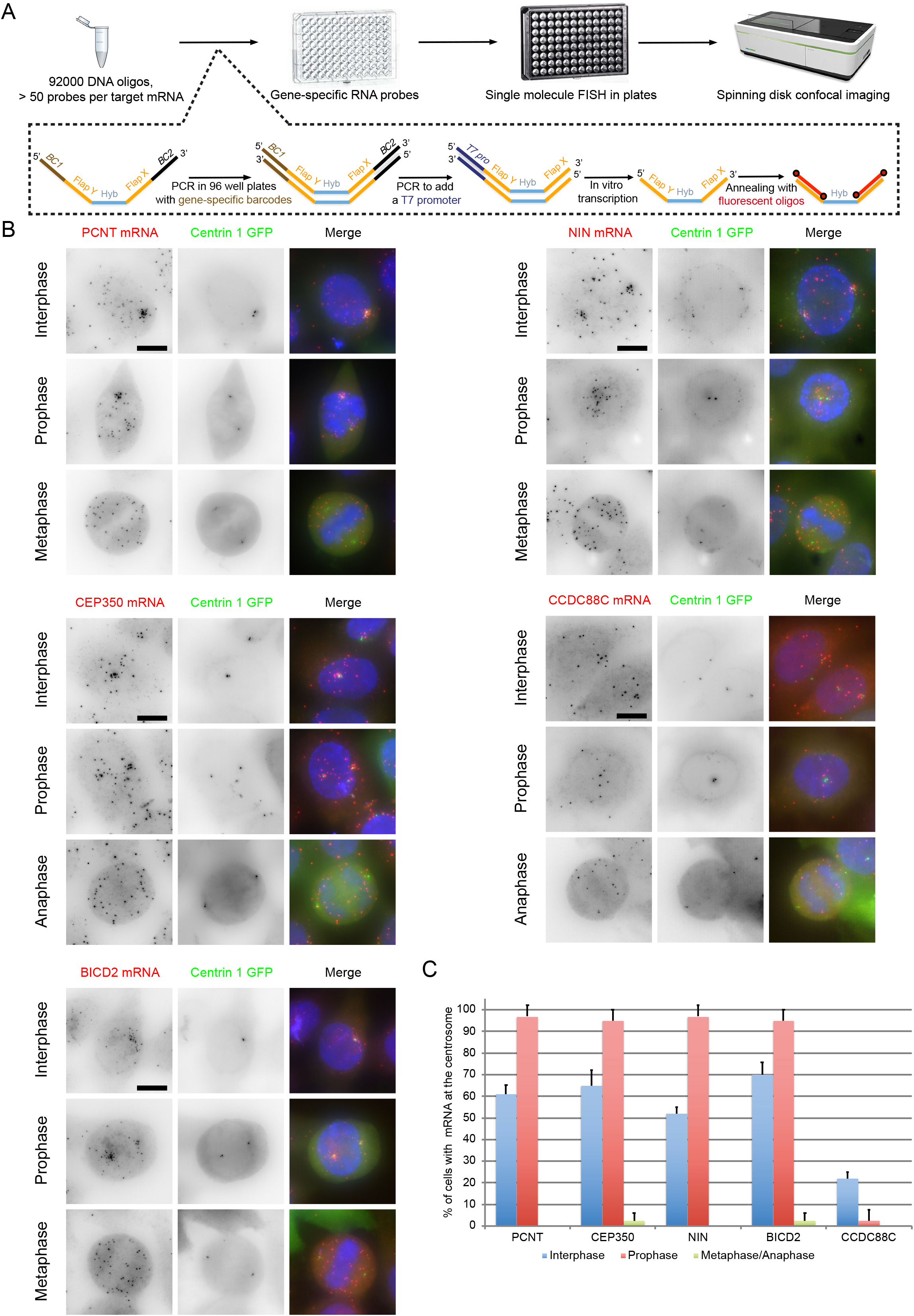
An mRNA screen identifies new mRNAs localizing to centrosomes. (A) Summary of the high-throughput smFISH pipeline. Top: starting from a pool of DNA oligonucleotides, gene-specific RNA probes are generated. SmFISH is performed on cells grown in 96 well plates, which are imaged by spinning disk confocal microscopy. Bottom: schematic representation of probe generation. BC: gene-specific barcode; FLAP X and Y: shared overhang sequences that hybridize with TYE-563 labeled LNA oligonucleotides (shown in red); Hyb: Hybridization sequence specific of the target mRNA, T7 pro: T7 RNA polymerase promoter. (B) Micrographs of HeLa cells stably expressing Centrin1-GFP and imaged by wide-field microscopy in interphase and mitosis. Left and red: Cy3 fluorescent signals corresponding to PCNT, CEP350, BICD2, NIN, or CCDC88C mRNAs labeled by low-throughput smiFISH; middle and green: fluorescent signals corresponding to the Centrin1-GFP protein. Blue: DNA stained with DAPI. Scale bars: 10 microns. (C) Histogram depicting the percentage of cells in each phase, showing the centrosomal localization of each mRNA (n=50 for interphase, and 25 for each mitotic phase; counts were performed twice from independent experiments). Data represent mean and standard deviation.

We screened a total of 602 genes using HeLa cells stably expressing a Centrin1-GFP fusion to label centrosomes. High-throughput spinning-disk microscopy was used to acquire full 3D images at high resolution (200 nm lateral and 600 nm axial), and two sets of images were recorded, to image either interphase or mitotic cells (see Materials and Methods). Centrosomal RNA enrichment was assessed by manual annotations of the images. These analyses yielded several localized mRNAs, including six that concentrated near centrosomes (Table 1 and Table S1). The localization of these mRNAs was then confirmed by performing low-throughput smiFISH. The results confirmed that the six candidate mRNAs accumulated at centrosomes during interphase and/or mitosis. These transcripts included PCNT and NUMA1 mRNA that were also recently identified by us and others (Sepulveda et al., 2018; Chouaib et al., 2019), as well as several new ones: NIN, BICD2, CCDC88C and CEP350 (Figure 1B, Table 1). Taking into account ASPM and HMMR that we also recently identified (Chouaib et al., 2019), a total of 8 mRNAs thus localize at centrosomes in human cells. These transcripts encode proteins that regulate various aspects of centrosome maturation, spindle positioning, and MT dynamics. Interestingly, the localization of these mRNAs varied during the cell cycle. CCDC88C mRNA localized during interphase but not mitosis. PCNT, NIN, BICD2, HMMR and CEP350 mRNAs localized during interphase and early mitosis, but delocalized at later mitotic stages (Figure 1C, and see below). In contrast, NUMA1 and ASPM mRNAs only localized during mitosis (Chouaib et al., 2019; see below). Since all these mRNA code for centrosomal proteins, centrosomes thus appear to have a dedicated translational program that is regulated during the cell cycle.

### ASPM, NUMA1 and HMMR proteins localize to centrosome at specific cell cycle stage

We then focused on ASPM, NUMA1 and HMMR, and analyzed in more detail their expression and localization. We first analyzed the expression of their respective proteins during the cell cycle. For this, we took advantage of HeLa Kyoto cell lines that stably express bacterial artificial chromosomes (BAC) containing the entire genomic sequences of the genes of interest, and carrying a C-terminal GFP tag (Poser et al., 2008; Chouaib et al., 2019). These BACs contain all the gene regulatory sequences and are expressed at near-endogenous levels and with the proper spatio-temporal pattern (Poser et al., 2008). Time-lapse microscopy of single cells revealed that ASPM-GFP and HMMR-GFP expression rose progressively during interphase to culminate just before mitosis, while that of NUMA1-GFP appeared constant during the cell cycle. Interestingly, ASPM-GFP and NUMA1-GFP had similar localization patterns. Both proteins were mainly nucleoplasmic during interphase and precisely initiated centrosomal localization in prophase. During cell division, they accumulated at the spindle pole with a weak labeling of the proximal spindle fibers (Figure S1A-B). In contrast, HMMR-GFP accumulated on the entire spindle throughout mitosis and furthermore concentrated on cytokinetic bridges in telophase. During interphase, it labeled microtubules (MTs) and localized to centrosomes several hours before cell division (Figure S1C).

### ASPM, NUMA1, and HMMR mRNAs localize to centrosome at the same time as their proteins

We next determined the localization of ASPM, NUMA1 and HMMR mRNAs during the different mitotic phases. We again used the GFP-tagged BAC HeLa cell lines to correlate protein and mRNA localization. smFISH was performed against the GFP RNA sequence using a set of 44 Cy3 labeled oligonucleotide probes. In interphase, ASPM-GFP and NUMA1-GFP mRNAs and proteins did not localize to centrosomes as previously reported (Chouaib et al., 2019), while HMMR-GFP mRNAs and its protein co-localized to centrosomes in a fraction of the cells. During mitosis, ASPM-GFP mRNAs and protein were enriched together on mitotic centrosomes across all phases of cell division (Figure 2A and 2B). In contrast, NUMA1 and HMMR mRNAs only accumulated at centrosomes during the early stages of cell division, prophase and prometaphase, where they co-localized with their protein. A random mRNA distribution was seen during metaphase and anaphase, although both proteins still remained on the mitotic spindle (Figure 2C and 2D). Unlike NUMA1, the centrosomal localization of HMMR mRNA and protein was re-established in telophase, where they accumulated together at the cytokinetic bridges (Figure 2E and 2F).

**Figure 2:**
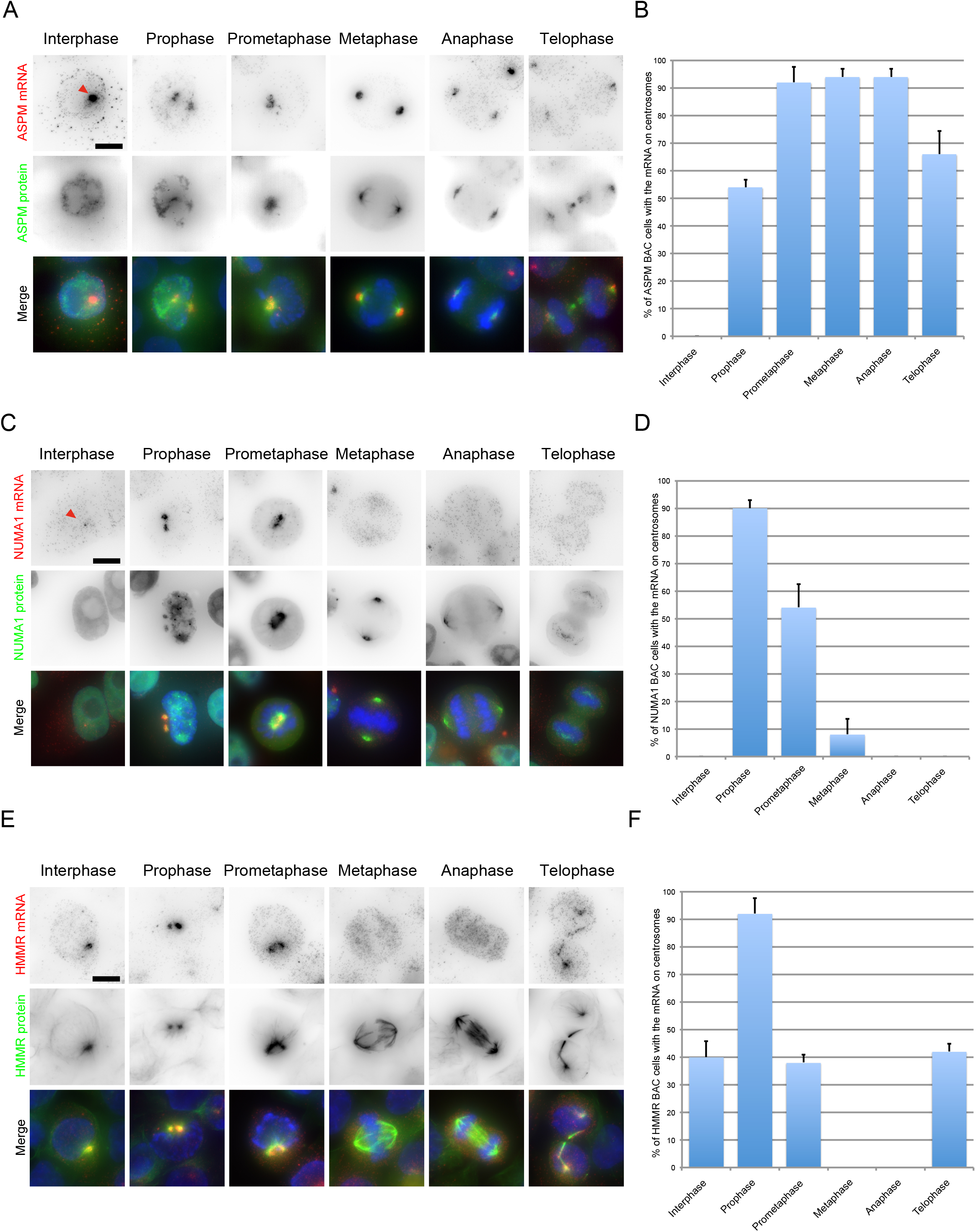
Differential centrosomal localization of ASPM, NUMA1 and HMMR mRNAs during mitosis. (A) Micrographs of HeLa cells expressing an ASPM-GFP BAC and imaged at different phases of the cell cycle. Top and red: Cy3 fluorescent signals corresponding to ASPM mRNAs labeled by smFISH; middle and green: GFP signals corresponding to the ASPM protein. Blue: DNA stained with DAPI. Scale bar: 10 microns. The red arrow indicates a transcription site. (B) Histogram depicting the percentage of cells in each phase showing centrosomal localization of ASPM mRNAs (n=25 cells per mitotic phase, repeated twice). Data represent mean and standard deviation. (C and D) Legend as in A and B, but for HeLa cells containing a NUMA1-GFP BAC. (E and F) Legend as in A and B, but for HeLa cells containing an HMMR-GFP BAC.

To detail these findings, we performed two-color smFISH experiments detecting one BAC-GFP mRNA in Cy3 and an endogenous mRNA in Cy5. We analyzed all pairwise combinations of ASPM, NUMA1 and HMMR mRNAs. To gain more precision, we divided each of prophase, prometaphase and telophase into two sub-phases, early and late (see Materials and Methods). During early prophase, NUMA1 and HMMR mRNAs could be seen on centrosomes but not ASPM mRNAs that only joined during late prophase (Figure S2). During early prometaphase, all three mRNAs were enriched on centrosomes. However, the centrosomal localization of NUMA1 and HMMR mRNAs became much less frequent starting at late prometaphase, while that of ASPM mRNA could still be observed in metaphase and anaphase (Figure S2 and S3). Finally, ASPM but not HMMR mRNAs accumulated on centrosomes during early telophase, while the opposite was observed at late telophase. Interestingly, the three mRNAs never perfectly co-localized on centrosomes at any of the cell-cycle stages: certain peri-centrosomal regions were occupied by one transcript while others contained the other mRNA (see Figures S2 and S3).

We recently showed that two translation factors, eIF4E and a phosphorylated form of RPS6, accumulated at centrosomes in prophase (Chouaib et al., 2019). Here, we examined their localization during the other mitotic phases. The centrosomal localization of these factors was strongest in prophase (unpublished data), and thus mirrored the bulk of centrosomal mRNAs, which, except for CCDC88C, localize to centrosome most strongly in prophase. The detection of translation factors together with mRNAs at centrosomes likely reflects where translation occurs. Taken together, these data demonstrate a fine tuning of spatio-temporal dynamics for the centrosomal localization of ASPM, NUMA1, and HMMR mRNAs, with each mRNA localizing at a specific stage and place during cell division.

### The localization of the 8 centrosomal mRNAs is inhibited by puromycin but not cycloheximide

Next, we analyzed the localization mechanism of these mRNAs and first questioned whether localization requires translation. To this end, we used a HeLa cell line expressing Centrin1-GFP to label centrosomes and treated it for 20 minutes with either cycloheximide, which blocks ribosome elongation, or puromycin, which induces premature chain termination. We first analyzed the mRNAs localizing during interphase (NIN, BICD2, CCDC88C, CEP350, HMMR and PCNT). Remarkably, these six mRNAs became delocalized after puromycin treatment while cycloheximide had no effect (Figure 3A and 3B). Long puromycin treatments prevent entry into mitosis. However, a 5-minute incubation was sufficient to inhibit the centrosomal localization of ASPM, NUMA1 and HMMR mRNAs at all the mitotic phases in which they normally localize (Figures S4, S5, and S6), while cycloheximide still had no effect. Since cycloheximide inhibits translation but leaves the nascent peptide chain on ribosomes, while puromycin removes it, our data suggest that mRNA localization to centrosomes requires the nascent peptide. RNA localization is thus expected to occur co-translationally for all the 8 mRNAs, pointing toward a common localization mechanism.

**Figure 3:**
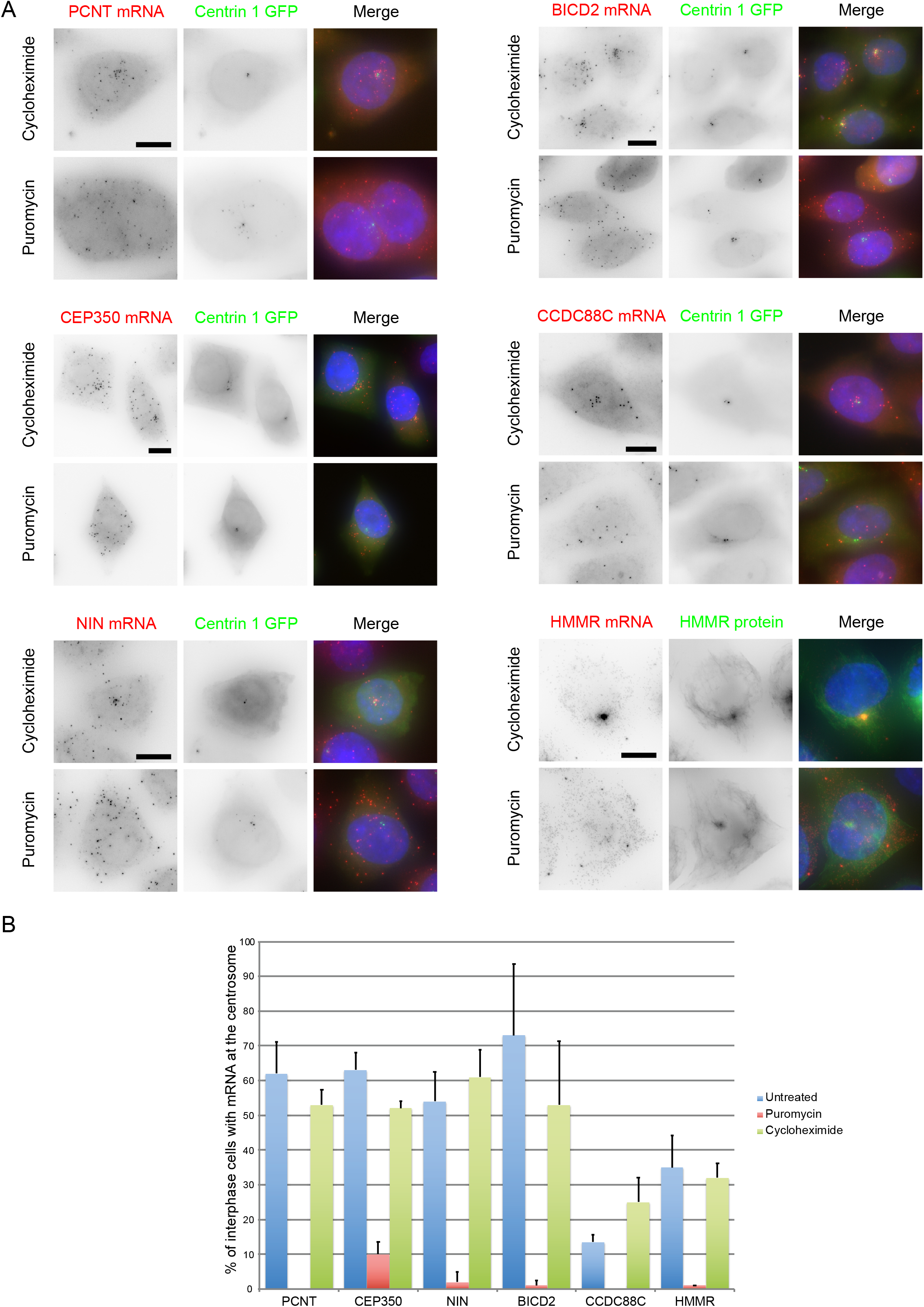
Translation initiation is required for the localization of the centrosomal mRNAs during interphase. (A) Micrographs of HeLa cells expressing either Centrin1-GFP or an HMMR-GFP BAC, and treated with cycloheximide or puromycin. Left and red: Cy3 fluorescent signals corresponding to PCNT, CEP350, BICD2, NIN, CCDC88C, or HMMR-GFP mRNAs labeled by smiFISH or smFISH; middle and green: GFP signals corresponding to either Centrin1-GFP or the HMMR protein. Blue: DNA stained with DAPI. Scale bars: 10 microns. (B) Histogram depicting the percentage of cells showing centrosomal localization of the mRNA after the indicated treatments (20 minutes; n=50 cells per condition, repeated twice). Data represent mean and standard deviation.

### Translation of ASPM coding sequence is necessary and sufficient for localizing its mRNA at centrosomes

To investigate how mRNAs localize to centrosomes in more detail, we focused on ASPM and first asked whether the 5’ and 3’ UTRs were necessary for its localization. To this end, a full-length ASPM mouse coding sequence (CDS) was fused to the C-terminal of GFP and expressed via transient transfection in HeLa Kyoto cells. To detect mRNAs produced from this reporter only, we performed smFISH with probes directed against the GFP RNA sequence. Mitotic cells expressing the plasmid could be identified by the accumulation of GFP-ASPM, which localized on centrosomes and the mitotic spindle. Interestingly, we could detect ASPM-GFP mRNAs on mitotic centrosomes in most of the transfected cells (Figure 4A). This demonstrated that the 5’ and 3’ UTRs of ASPM mRNA are not required for its centrosomal enrichment.

**Figure 4:**
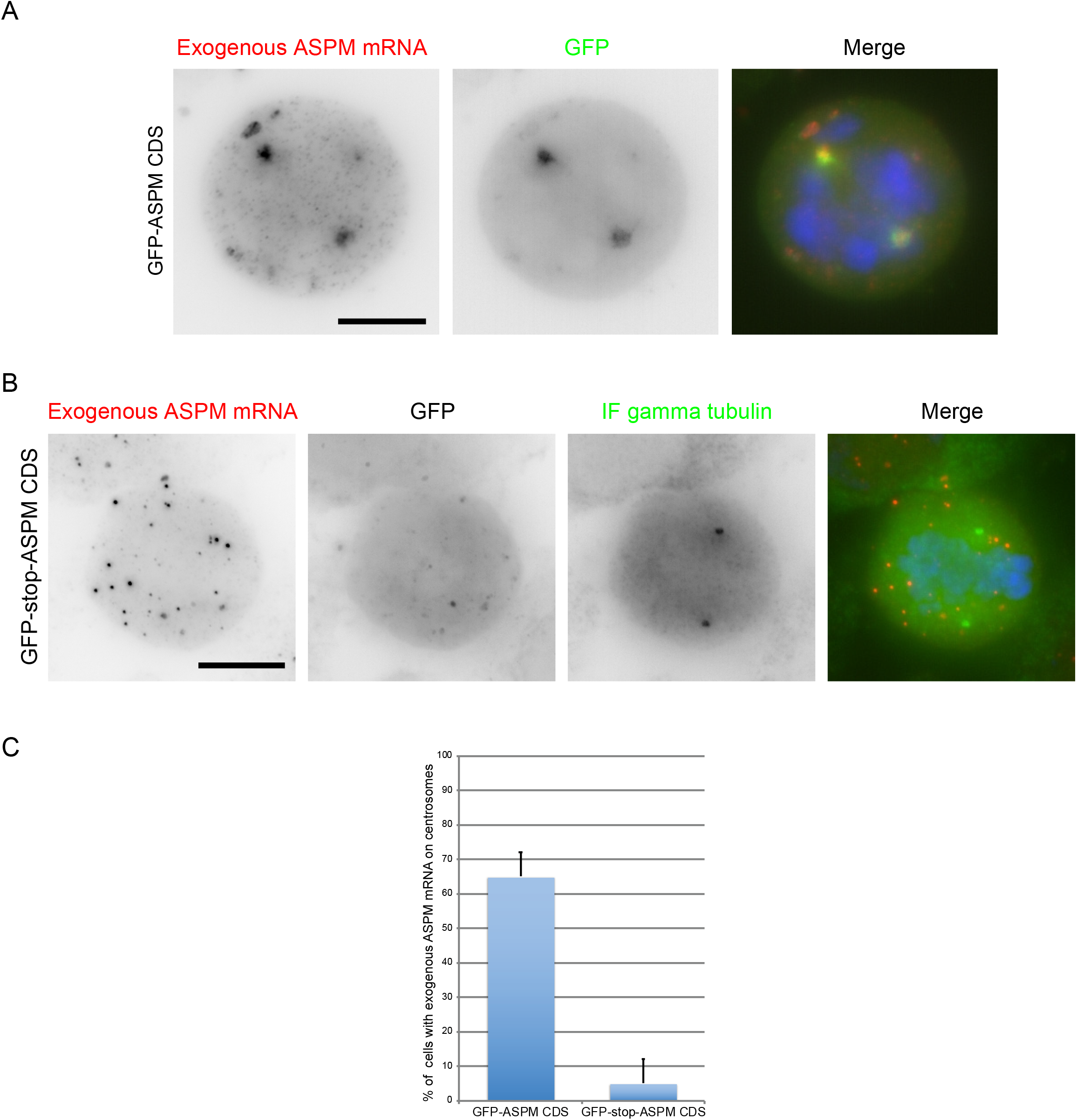
Translation of the ASPM cDNA is necessary and sufficient for its centrosomal localization. (A) Micrographs of mitotic HeLa cells transiently expressing GFP-ASPM CDS. Left and red: Cy3 fluorescent signals corresponding to the exogenous mRNAs labeled by smFISH; middle and green: GFP signals corresponding to the exogenous proteins. Blue: DNA stained with DAPI. Scale bar: 10 microns. (B) Legend as in A, but with cells expressing GFP-stop-ASPM CDS and with an IF against gamma-tubulin. DNA is shown in blue, centrosomes in green and the mRNA in red. (C) Histogram depicting the percentage of mitotic cells expressing each construct and showing centrosomal localization of exogenous ASPM mRNAs (n=20 cells, repeated twice). Data represent mean and standard deviation.

Next, we explored how the same GFP-ASPM mRNA would localize if the nascent ASPM protein was not translated. To test this, we introduced a stop codon between the GFP and ASPM coding sequences, generating a GFP-stop-ASPM construct. Transient transfection showed a diffuse GFP signal as expected. Interestingly, mRNAs translating this reporter failed to localize to mitotic centrosomes labeled by an immunofluorescence (IF) against gamma tubulin (Figure 4B and 4C). Taken together, this demonstrated that the nascent ASPM polypeptide is required for trafficking its own transcript to mitotic centrosomes.

### ASPM mRNAs are actively transported toward centrosomes and anchored on the mitotic spindle

To gain more insights into the localization mechanism, we imaged the endogenous ASPM mRNAs in living mitotic cells. To this end, we inserted 24 MS2 repeats in the 3’ UTR of the endogenous gene, using CRISPR/Cas9 mediated homology-directed repair in HeLa Kyoto cells (Figure 5A). Heterozygous clones were confirmed by genotyping (Figure S7A). Moreover, two-color smFISH performed with MS2 and ASPM probes showed that the tagged mRNA accumulated at centrosomes in mitosis (Figure 5B), indicating that the MS2 sequences did not interfere with localization. We then stably expressed low levels of the MS2-coat protein (MCP) fused to GFP and a nuclear localization signal (MCP-GFP-NLS). This fusion protein binds the MS2 repeat and allows to visualize the tagged RNA in living cells (Bertrand et al., 1998). Indeed, mitotic cells expressing MCP-GFP-NLS displayed diffraction limited fluorescent spots that localized near the centrosomes. Moreover, these spots co-localized with single RNA molecules revealed with probes against either endogenous ASPM mRNA or the MS2 tag, indicating that binding of MCP-GFP-NLS to the tagged mRNA did not abolish RNA localization to the centrosome (Figure S7B).

**Figure 5:**
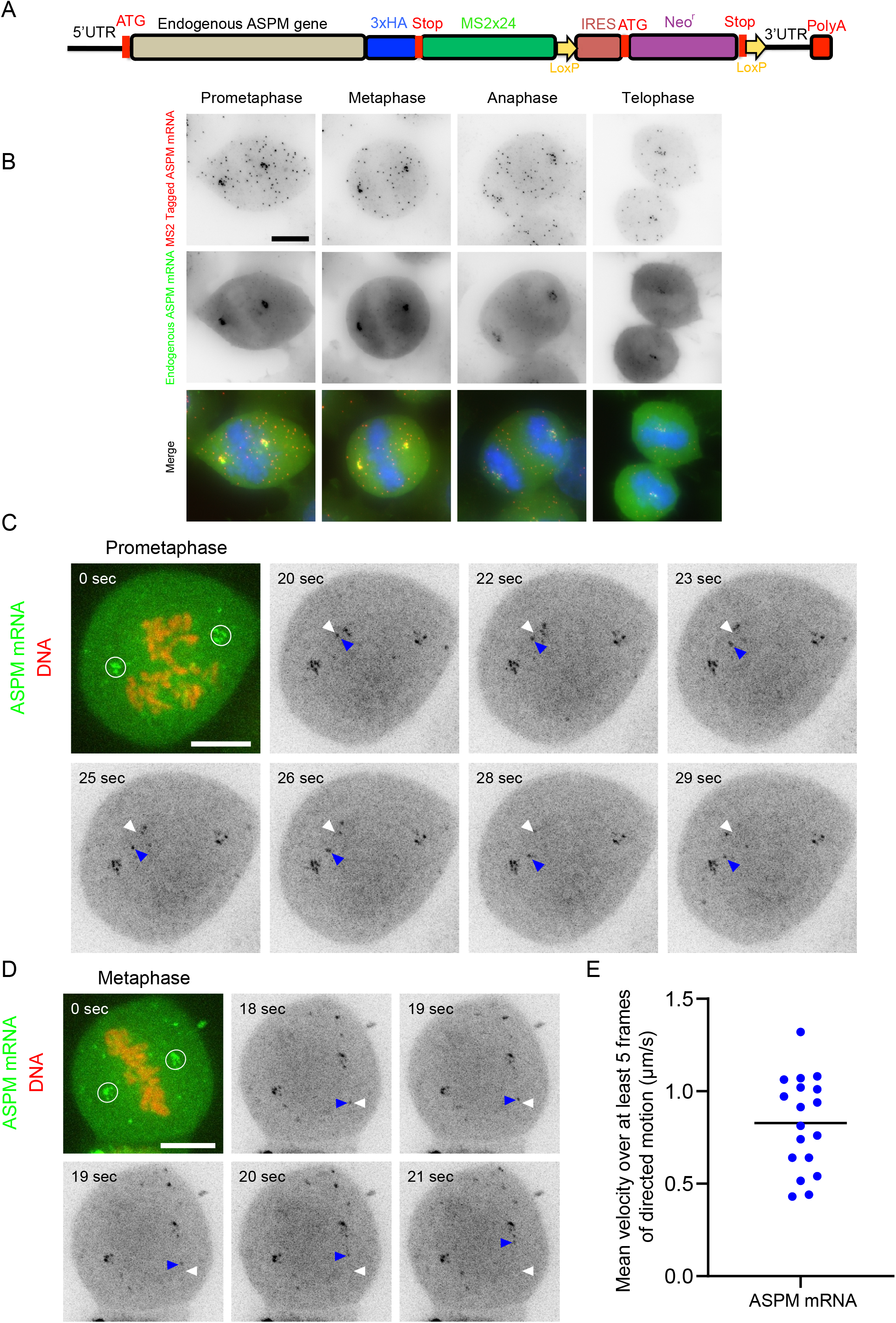
ASPM mRNAs undergo diffusion, directed movements, and anchoring to spindle poles and fibers across mitosis. (A) Schematic representation showing the insertion of 24 MS2 repeats at the end of the ASPM gene using CRISPR-Cas9. HA: human influenza hemagglutinin tag. Stop: stop codon. IRES: internal ribosome entry site. Neo^r^: neomycin resistance. UTR: untranslated region. PolyA: poly A tail. (B) Images are micrographs of mitotic HeLa cells with an ASPM-MS2×24 allele. Upper and red: Cy3 fluorescent signals corresponding to tagged mRNAs labeled by smFISH with MS2 probes; middle and green: Cy5 signals corresponding to the tagged and untagged ASPM mRNA detected by smiFISH with endogenous ASPM probes. Blue: DNA stained with DAPI. Scale bar: 10 microns. (C) Snapshots of ASPM-MS2×24 cells expressing MCP-GFP-NLS and imaged live during prometaphase. In the first panel, the GFP signal is shown in green and corresponds to ASPM mRNAs labeled by the MS2-MCP-GFP-NLS. The Cy5 signal is shown in red and corresponds to DNA. Scale bar: 10 microns. Time is in seconds. The white circle displays ASPM mRNAs with limited diffusion. In subsequent panels, only the GFP signal is shown and is represented in black. White arrowheads indicate the starting position of an mRNA molecule, while blue ones follow its current position. (D) Legend as in (C) but for a cell captured in metaphase. (E) Dot plot showing the speed of ASPM mRNA molecules displaying directed motion, calculated over at least 5 frames (or 2.5 seconds). The horizontal line represents the mean.

Next, we performed live cell experiments. We labeled DNA using SiR-DNA to identify the mitotic phase and we imaged cells in 3D at a rate of 2-4 fps using spinning disk microscopy. During prometaphase, three populations of ASPM mRNA molecules were observed: (i) mRNAs diffusing in the cytosolic space, (ii) immobile molecules that corresponded to mRNAs localizing at the centrosome, and (iii) mRNAs undergoing directed movements towards the centrosome (Figure 5C). The thickness of mitotic cells yielded low signal-to-noise ratios which, combined with the rapid movements of the mRNAs, made single particle tracking with automated software difficult. We thus manually tracked mRNA molecules undergoing directed movements and observed that they moved at speeds ranging from 0.5 to 1 μm/sec (Figure 5E), which is compatible with motor-mediated transport (Schiavo et al., 2013).

In metaphase and anaphase where the mRNA is at centrosomes, we expected several possibilities for the movement of mRNA molecules: (i) stable anchoring to the centrosome; (ii) diffusion within a confined space around the centrosome; or (iii) diffusion away from centrosomes and re-localization by a motor-dependent mechanism. Live imaging revealed that ASPM mRNA localizing at mitotic centrosome did not diffuse and were immobile (Figure 5D, Movie 1). In addition, directed movements toward centrosomes were also observed in mid-mitosis albeit to a lesser extent than in prometaphase (Figure 5D arrows, Movie 1). Interestingly, we also observed that some ASPM mRNAs were attached on the spindle fibers rather than on the spindle poles (Movie 2). Taken together, these live imaging experiments demonstrated that ASPM mRNAs are actively transported to the mitotic centrosomes at the onset of mitosis and are then anchored on the spindle poles and fibers.

### ASPM polysomes are actively transported towards centrosomes during prophase and prometaphase

Since the localization of ASPM transcripts required in-*cis* translation and might thus involve the transport of translated mRNAs, we next imaged ASPM polysomes. To this end, we used the SunTag system that allows to image nascent polypeptide chains (Morisaki et al., 2016; Pichon et al., 2016; Wang et al., 2016; Wu et al., 2016; Yan et al., 2016). The SunTag is composed of a repeated epitope inserted in the protein of interest and a singlechain antibody fused to GFP. Binding of the fluorescent antibody to the epitope occurs when it emerges from the ribosome, and this allows visualizing nascent protein chains and polysomes in live cells. We engineered a HeLa Kyoto cell line with 32 SunTag repeats fused to the 5’ end of the ASPM gene, using CRISPR/Cas9 mediated homology-directed repair (Figure 6A). Heterozygous clones were confirmed by genotyping (Figure S8A). The cells were then transduced with a lentivirus expressing the scFv monochain antibody fused to GFP (scFv-sfGFP). Bright GFP foci were observed and confirmed to be polysomes based on both their sensitivity to puromycin (Figure S8B) and co-localization with endogenous ASPM mRNA by smiFISH (Figure 6B). Moreover, ASPM mRNAs having the SunTag accumulated on mitotic centrosomes (Figure S8C), indicating that the tagging process did not abolish centrosomal RNA targeting.

**Figure 6:**
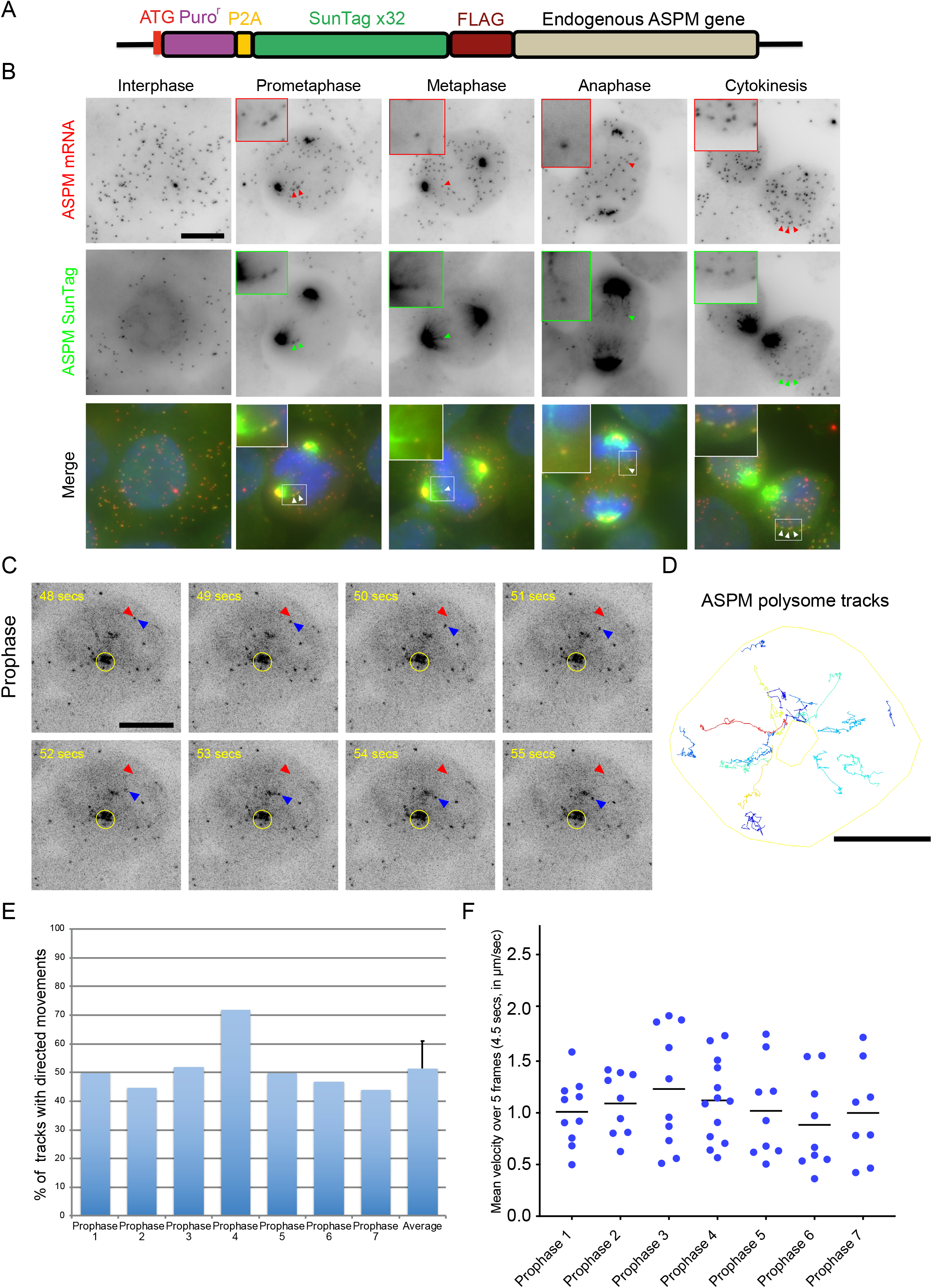
ASPM polysomes show directed movements towards the centrosome in early mitosis. (A) Schematic representation of a cassette containing 32 SunTag repeats and with recombination arms to target insertion at the N-terminus of the endogenous ASPM gene. Puro^r^: puromycin resistance; P2A: self-cleaving signal; FLAG: octapeptide FLAG tag. (B) Micrographs of a HeLa cell with a SunTagx32-ASPM allele and expressing scFv-sfGFP, imaged during interphase and mitosis. Upper and red: Cy3 fluorescent signals corresponding to tagged and untagged ASPM mRNAs labeled by smiFISH with probes against the endogenous ASPM mRNA; middle and green: GFP signals corresponding to ASPM polysomes and mature protein. Blue: DNA stained with DAPI. Scale bar: 10 microns. Red and green arrowheads indicate ASPM mRNAs and polysomes, respectively. White arrows indicate the overlay of red and green arrows. Insets represent zooms of the white-boxed areas. (C) Snapshots of HeLa cells with a SunTagx32-ASPM allele and expressing scFv-sfGFP, imaged live during prophase. The SunTag signal is shown in black and corresponds to ASPM polysomes and mature proteins. Scale bar: 10 microns. Time is in seconds. The yellow circle indicates ASPM mature protein. Red arrowheads indicate the starting position of an ASPM polysome, while blue ones follow its position at the indicated time. (D) A TrackMate overlay of the same cell in C showing polysomes tracks. Color code represents displacement (dark blue lowest, red highest). The outer yellow outline represents the cell border, while the inner one represents centrosomes marked by the mature ASPM protein. Scale bar: 10 microns. (E) A histogram showing the percentage of polysomes displaying directed movements towards the centrosome during prophase. The last column represents the mean and standard deviation. (F) Dot plot showing the average speed of ASPM polysomes displaying directed motions (calculated over at least 5 frames), in prophase cells. The horizontal line represents the mean.

We first imaged SunTag-ASPM polysomes and mRNA together in fixed cells (Figure 6B). In early prophase, most mRNAs and polysomes did not localize to centrosomes, in agreement with the smFISH data (Figure S2). In prometaphase, metaphase and anaphase, the accumulation of the SunTag-ASPM mature protein at the spindle poles prevented visualizing ASPM polysomes at this location. However, some ASPM polysomes were observed outside the spindle area indicating that translation was pursued during the entirety of mitosis (Figure 6B). At the end of telophase, ASPM mRNAs could be seen translated on the nuclear envelope as previously reported for cells in interphase (Chouaib et al., 2019).

We then performed live imaging in 3D at acquisition rates of 1-1.3 stacks per second, using spinning disk microscopy. We first imaged cells in prophase (Movie 3). Remarkably, while many ASPM polysomes were dispersed in the cytoplasm at the beginning of prophase, they displayed rapid directed motions towards the centrosome, leading to their accumulation at this location at the end of the movie (Figure 6C, Movie 3). Single particle tracking showed that an average of 51% of ASPM polysomes displayed such directed movements in early prophase (Figure 6D, E). Directed movements were also detected in prometaphase but less frequently (Movie 4). Calculating the average velocity of polysomes undergoing directed movements showed that their speed ranged from around 0.25 to 2 μm/s, which is compatible with the velocities of both ASPM mRNAs and motor-dependent transport (Figure 6F). Taken together, this data directly proved that ASPM polysomes were actively transported to the centrosome at the onset of mitosis.

### ASPM mRNAs are translated on MTs during interphase

Similar to MS2-tagged ASPM mRNAs, some SunTag-ASPM polysomes did not co-localize with the spindle poles but rather with spindle fibers that were weakly labeled by the mature SunTag-ASPM protein (Figure 6B, arrows). This prompted us to investigate in more details the role of microtubules in the metabolism of ASPM mRNAs. We labeled MTs in living cells using a far-red dye (SiR-Tubulin) and performed sequential two-color 3D live imaging using a spinning disk microscope. We first imaged interphase cells and remarkably, we observed that many ASPM polysomes remained stably anchored to MTs during the course of the movies (66%; Figure 7A-C, Figure S9A, and Movie 5). In addition, we also observed directed motion of single polysomes, albeit at a low frequency (around 4%). We then characterized in more details the movements of ASPM polysomes and for this we classified them in four categories: (i) polysomes localizing on MTs; (ii) localizing at the nuclear envelope (as previously reported; Chouaib et al., 2019); (iii) niether localizing on MTs nor at the nuclear envelope, and thus free in the cytosol; and (iv) showing directed transport (Figure 7C, Figure S9A). The histogram of displacements between consecutive frames revealed a diffusion coefficient of 0.011μm^2^/s for polysomes on MTs, 0.004 μm^2^/s for the ones on the nuclear envelope and 0.041 μm^2^/s for those not on MT or the envelope (i.e. freely diffusing; Figure S9 B-D). We also calculated the mean square displacement (MSD) as a function of time (Figure 7D). This confirmed that the ASPM polysomes that were free in the cytosol diffused several folds faster than those bound to MTs or the nuclear envelope. As a control, we depolymerized MTs with a 10-minute nocodazole treatment before starting imaging. SiR-tubulin labeling confirmed the absence of MTs in treated cells, and we then tracked ASPM polysomes, excluding the ones attached to nuclear envelope (Figure S10A-C, Movie 6). The histogram of displacements and MSD curve revealed a single population with a diffusion coefficient of 0.035μm^2^/s, similar to polysomes not on MT or the nuclear envelope in untreated cells (Figure 7D and Figure S9E). Overall, this showed that a large fraction of ASPM mRNAs are locally translated on MTs during interphase, and that these polysomes are stably anchored to MTs, thereby limiting their diffusion.

**Figure 7:**
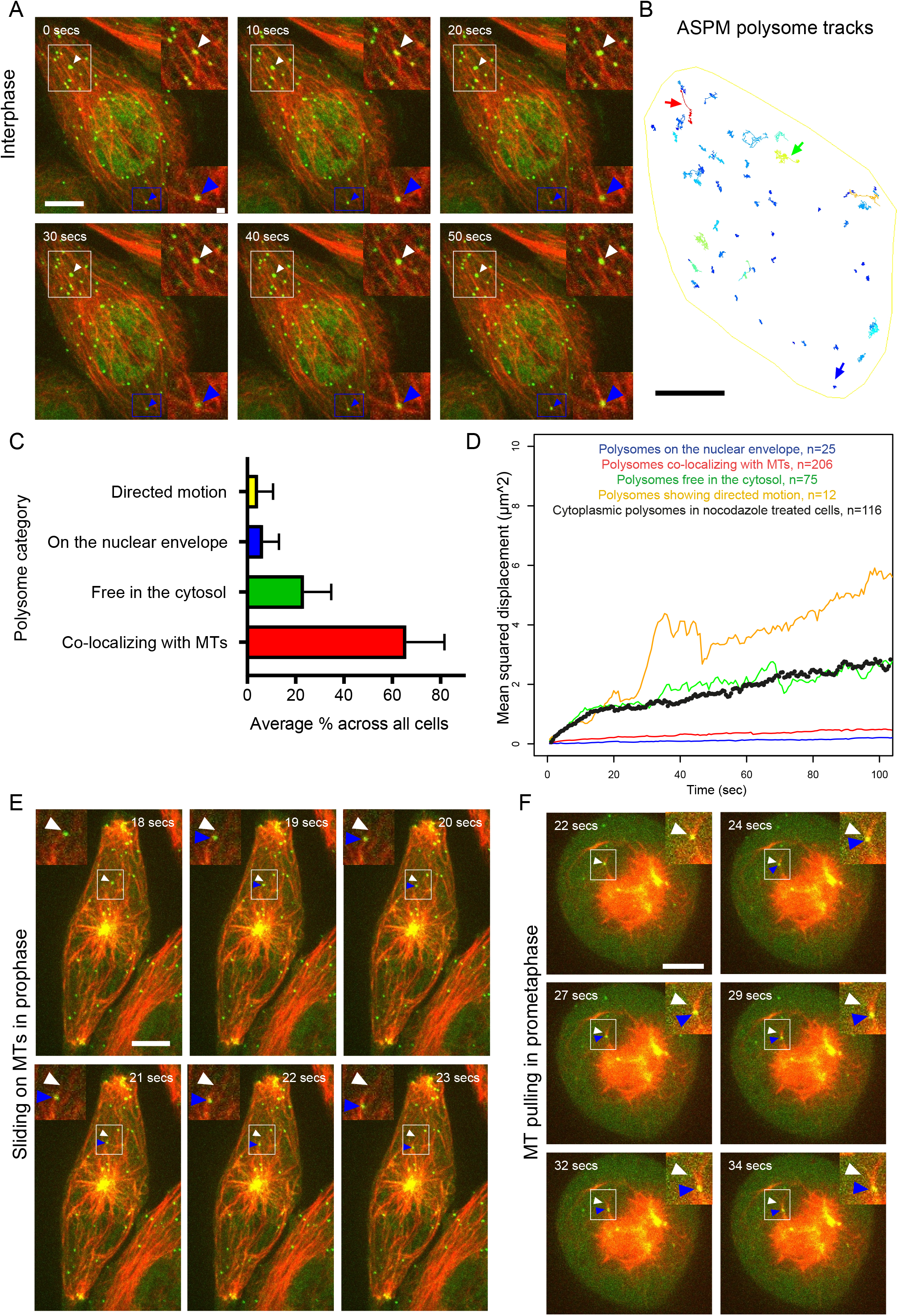
ASPM polysomes are anchored to microtubules in interphase and transported towards the centrosome in early mitosis. (A) Snapshots of SunTagx32-ASPM cells expressing scFv-sfGFP and imaged live during interphase with MT labeling. The SunTag signal is shown in green and corresponds to ASPM polysomes and mature proteins; the far-red signal is shown in red and corresponds to a tubulin staining. Scale bar: 10 microns. Time is in seconds. Upper and lower insets represent one-plane zooms of the white- and blue-boxed areas respectively. Arrowheads point to ASPM polysomes. (B) A TrackMate overlay of the same cell as in A showing the polysome tracks. Color code represents displacement (dark blue lowest, red highest). Scale bar: 10 microns. The red arrow indicates a track showing directed movement, the green one is a track that does not localize with MTs, and the orange one is a track that localizes with MTs. (C) The average percentage of ASPM tracks grouped in four categories depending on subcellular localization and behavior in interphase. Polysomes free in the cytosol: polysomes not on MT and not on the nuclear envelope. (D) Graph showing the mean squared displacement (MSD, in micron^2^) of polysomes as a function of time (in seconds) for each of the indicated polysome category and after a nocodazole treatment. Polysomes free in the cytosol: polysomes not on MT and not on the nuclear envelope. (E) Snapshots of SunTagx32-ASPM cells expressing the scFv-sfGFP and imaged live during prophase with MT labeling. The SunTag signal is shown in green and corresponds to ASPM polysomes and mature proteins; the far-red signal is shown in red and corresponds to a tubulin staining. Scale bar: 10 microns. Time is in seconds. Insets represent one-plane zooms of the white-boxed areas. White and blue arrowheads follow the initial and current position of a polysome, respectively. (F) Same legend as in E, but during prometaphase.

### ASPM polysomes are transported to mitotic centrosomes by either sliding on MTs or being pulled with entire MTs

To assess whether MTs are necessary for ASPM mRNA localization to centrosomes, we combined a brief nocodazole treatment (10 minutes) with smFISH, using the ASPM SunTag clone. A 10-minute treatment depolymerized MTs whereas centrosomes were still visible (Figure S10A, white arrowheads). ASPM mRNAs and polysomes no longer accumulated at mitotic centrosomes after MT depolymerization, despite the fact that the mRNAs were still translated (Figure S10 D-F). This indicated that intact MTs are required for ASPM mRNA localization.

We then performed dual color imaging of MTs and ASPM polysomes during mitosis. Tracks were shorter than in the previous mono-color movies because maintaining high framerates required the recording of only 3 Z planes in two-color experiments, as opposed to 15-20 in the single-color movies. Nevertheless, this allowed us to distinguish two types of movements towards centrosomes. In the first, ASPM polysomes rapidly slided along an immobile MT (Figure 7E, Movie 7). This likely corresponded to motor-driven movements of polysomes along MT cables. In the second type of movements, an ASPM polysome is stably attached to a MT and both were pulled together towards the centrosome (Figure 7F, Movie 8). In this case, the MT appears being hauled towards the centrosome and drags a tethered polysome with it. Indeed, MTs pulling and sliding has been previously described during mitosis (Forth and Kapoor, 2017; Enos et al., 2018). This demonstrated that ASPM polysomes are transported to the mitotic centrosomes via two mechanisms: sliding on MT, and tethering to MT coupled to MT remodeling.

### NUMA1 mRNAs and polysomes also display directed transport towards the centrosome in early mitosis

To assess the generality of the mechanism found with ASPM, we tagged another centrosomal mRNA, NUMA1. Using CRISPR/Cas9 mediated homology-directed repair in HeLa Kyoto cells, we generated a clone with an MS2×24 tag in the NUMA1 3’UTR, and another clone with a SunTag fused to the N-terminus of the protein. Proper recombination was verified by genotyping (Figure S11 A, B). In the NUMA1 MS2×24 clone, two-color smFISH performed against either the MS2 tag or the endogenous NUMA1 mRNAs revealed that MS2 tagging did not prevent NUMA1 mRNA from localizing to mitotic centrosomes (Figure 8A). Likewise, the SunTagged NUMA1 mRNA also localized to mitotic centrosomes (Figure S11C). Furthermore, in the SunTag NUMA1 clone, smiFISH against the endogenous NUMA1 mRNA revealed that the mRNA colocalized with bright SunTag foci in both interphase and mitosis (Figure 8B and S11D). A puromycin treatment removed these bright cytoplasmic SunTag foci, confirming that they were NUMA1 polysomes (Figure S11E). Therefore, tagging the endogenous NUMA1 mRNA with MS2 and SunTag repeats did not abolish its centrosomal localization.

**Figure 8:**
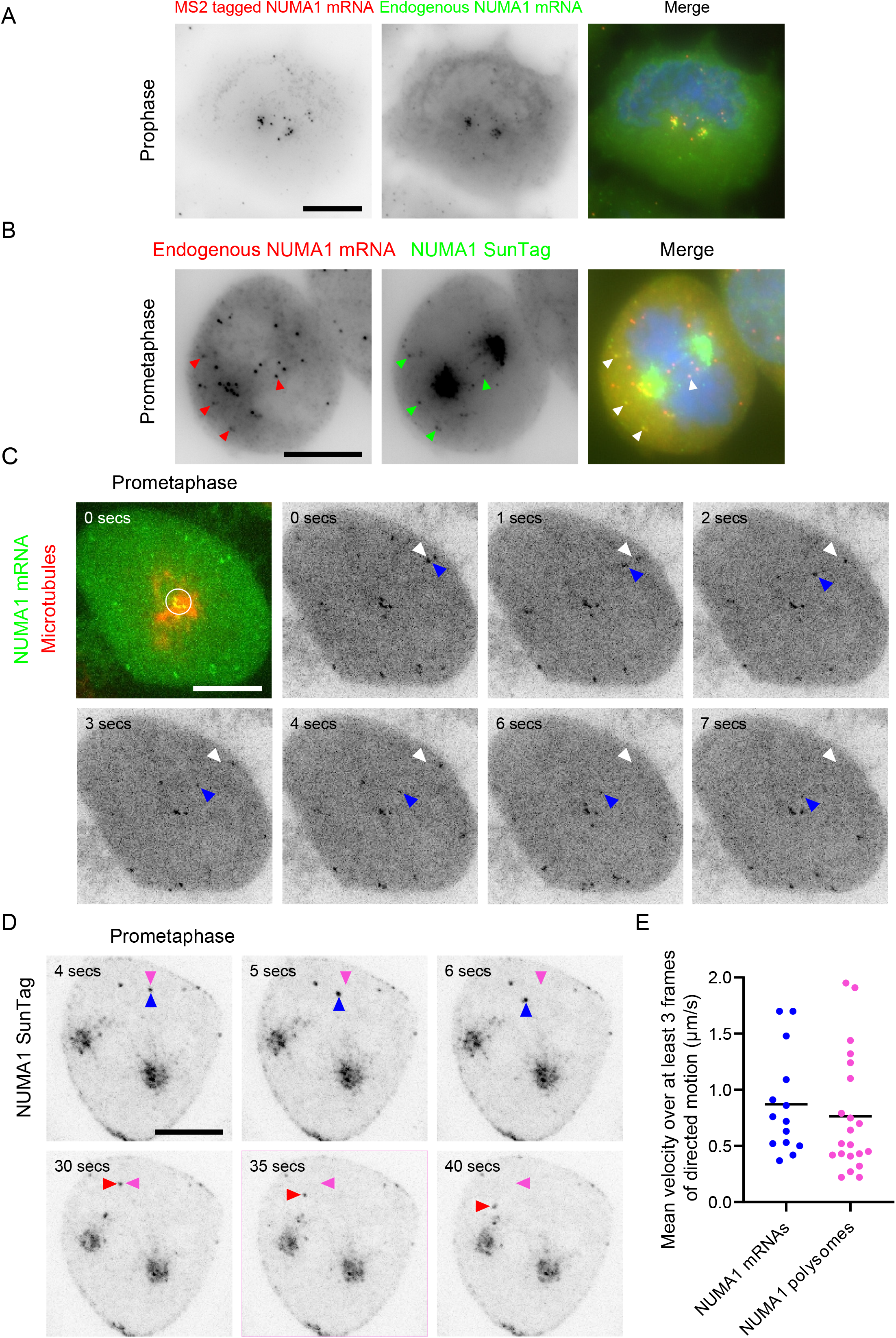
NUMA1 mRNAs and polysomes also show directed movements towards the centrosome in early mitosis. (A) Images are micrographs of HeLa cells with an NUMA1-MS2×24 allele and imaged in prophase. Left and red: Cy3 fluorescent signals corresponding to tagged mRNAs labeled by smFISH with MS2 probes; middle and green: Cy5 signals corresponding to the tagged and untagged NUMA1 mRNA labeled by smiFISH with probes against the endogenous mRNA. Blue: DNA stained with DAPI. Scale bar: 10 microns. (B) Images are micrographs of SunTagx32-NUMA1 cells expressing the scFv-sfGFP and imaged during prometaphase. Left and red: Cy3 fluorescent signals corresponding to NUMA1 mRNAs labeled by smiFISH; middle and green: GFP signals corresponding to NUMA1 polysomes and mature protein. Blue: DNA stained with DAPI. Scale bar: 10 microns. Red and green arrowheads indicate NUMA1 mRNAs and polysomes, respectively. White arrows indicate the overlay of red and green arrows. (C) Snapshots of NUMA1-MS2×24 cells expressing MCP-GFP-NLS and imaged live during prometaphase. In the first panel, the GFP signal is shown in green and corresponds to NUMA1 mRNAs labeled by the MS2-MCP-GFP-NLS. The far-red signal is shown in red and corresponds to microtubules. Scale bar: 10 microns. Time is in seconds. The white circle indicates NUMA1 mRNAs with limited diffusion. In subsequent panels, only the GFP signal is shown and is represented in black. White arrowheads indicate the starting position of an mRNA molecule, while blue ones follow its current position. (D) Snapshots of SunTagx32-NUMA1 cells expressing the scFv-sfGFP and imaged live during prometaphase. The SunTag signal is shown in black and corresponds to NUMA1 polysomes and mature proteins. Scale bar: 10 microns. Time is in seconds. Pink arrowheads indicate the starting position of two polysomes, while blue and red ones follow its current position. (E) Dot plot showing the average speeds of NUMA1 mRNAs and polysomes displaying directed motion (calculated over at least 3 frames). The horizontal line represents the mean.

Following stable expression of MCP-GFP-NLS or scFv-sfGFP, we imaged single NUMA1 mRNAs and polysomes, respectively, in living interphase and mitotic cells. In interphase, we could observe some NUMA1 polysomes undergoing rapid rectilinear movements (Figure S11F). Remarkably, in prometaphase, we observed rapid directed motion of both mRNAs and polysomes toward the centrosome, indicating that NUMA1 polysomes are actively transported towards centrosomes at the onset of mitosis, similar to what we observed for ASPM (Figure 8C-D, Movies 9 and 10). We manually calculated the mean particle velocity. Both NUMA1 mRNAs and polysomes had average speeds of around 0.9 μm/s, which is compatible with motor directed movements and also similar to the velocities measured for ASPM (Figure 8E). These live imaging experiments of endogenous transcripts and polysomes show that active polysome transport is a localization mechanism shared by several centrosomal mRNAs.

### Drosophila orthologs of the human centrosomal mRNAs also localize to centrosomes and also require the nascent protein

We next examined whether centrosomal mRNA localization is evolutionary conserved. To this end, we investigated the localization of the *Drosophila* orthologs of the human centrosomal mRNAs. Out of the 8 human mRNAs, 5 had clear orthologs: ASPM (Asp), NUMA1 (Mud), BICD2 (BicD), CCDC88C (Girdin), and PCNT (Plp). We used S2R+ cells as a model and co-labeled these mRNAs with centrosomes, using smFISH coupled to IF against gamma-tubulin. Remarkably, we observed centrosomal enrichment for 4 of these 5 *Drosophila* mRNAs (Figure 9, Table 1). Moreover, while we could not obtain clear signals for the fifth mRNA in S2R+ cells (Plp), it was annotated as centrosomal in a previous large-scale screen (see Table 1). Interestingly, we also observed a cell-cycle dependent localization for Asp, Mud, and Girdin mRNAs (Figure S12). A large fraction of cells in interphase did not localize Mud mRNAs whereas it localized to centrosomes during mitosis (metaphase, anaphase, and telophase; Figure 9B and S12). Asp mRNA showed a similar dynamic, with the exception that it was not localized in telophase. In contrast, Girdin mRNAs only accumulated on mitotic centrosomes during telophase (Figure S12). These observations indicate that a conserved cell-cycle regulated centrosomal translational program occurs in both human and *Drosophila* cells.

**Figure 9:**
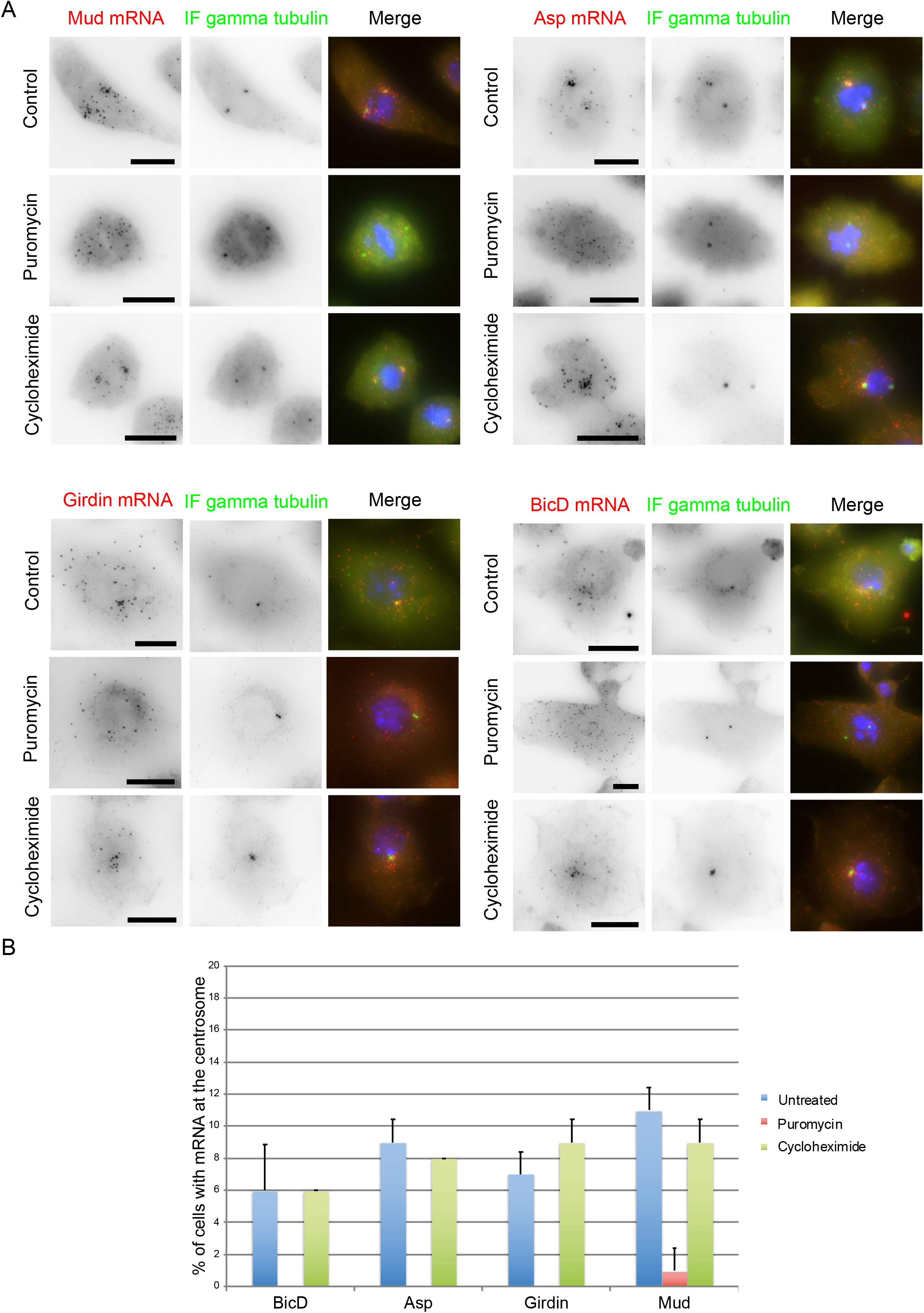
Translation-dependent targeting of centrosomal mRNA is conserved in *Drosophila*. (A) Images are micrographs of S2R+ cells treated with puromycin, cycloheximide or untreated as controls. Left and red: Cy3 fluorescent signals corresponding to centrosomal mRNAs labeled by smiFISH; middle and green: GFP signals corresponding to the gamma tubulin protein revealed by IF. Blue: DNA stained with DAPI. Scale bar: 10 microns. (B) Histogram depicting the percentage of cells in each condition showing centrosomal mRNA localization (n=50 cells, repeated twice). Data represent the mean and standard deviation.

Finally, we tested whether mRNAs in *Drosophila* depend on their nascent peptide to localize to centrosomes. We performed puromycin and cycloheximide treatments and co-labeled the mRNAs and centrosomes in S2R+ cells. As in human cells, puromycin treatment abolished centrosomal accumulation of all 4 mRNAs while cycloheximide did not (Figure 9). This indicated that not only the identity of centrosomal mRNAs is conserved from human to *Drosophila,* but also the localization mechanism, which requires the nascent polypeptide in both cases.

## Discussion

Here, we studied centrosomal mRNA localization in human cells. We uncovered a complex choreography of mRNA trafficking at centrosomes, particularly during mitosis, and we provide definitive evidence for a nascent chain-dependent transport of polysomes by motors and MTs. Remarkably, both the identity of localized mRNAs and the localization mechanism appear conserved from *Drosophila* to humans.

### A targeted smFISH screen reveals that centrosomal mRNAs are conserved from human to *Drosophila*

We used high-throughput smFISH to screen 602 genes encoding nearly the entire centrosomal proteome and we identified four new mRNAs localized at the centrosome (Table 1; Table S1). In total, eight human mRNAs now belong to this class: NIN, CEP350, PCNT, BICD2, CCDC88C, ASPM, NUMA1 and HMMR. All the corresponding proteins localize to the centrosome suggesting that their mRNAs are locally translated. Most of them also perform important centrosomal functions. NIN is localized to the sub-distal appendage of mother centrosomes and it functions in microtubule nucleation as well as centrosome maturation (Ou et al., 2002; Stillwell et al., 2004). CEP350 is also localized to sub-distal appendages and it is important for centriole assembly and MT anchoring to centrosomes (Yan et al., 2006; Le Clech, 2008). Pericentrin (PCNT) is a major component of the pericentriolar material (PCM) and it plays a structural role by bridging the centrioles to the PCM (Lee and Rhee, 2011). BICD2 contributes to centrosomal positioning and to centrosomal separation at the onset of mitosis (Raaijmakers et al., 2012; Splinter et al., 2010). ASPM and NUMA1 are two MT minus end binding proteins that accumulate at centrosomes during mitosis, and they control several aspects of spindle assembly and function (Jiang et al., 2017; Seldin et al., 2016). Finally, HMMR acts to separate centrosomes and to nucleate MT during spindle assembly. It also modulate the cortical localization of NUMA-dynein complexes to correct mispositioned spindles (Connell et al., 2017). The diversity of functions performed by these proteins suggests that RNA localization and local translation play an important role for the centrosome.

Remarkably, we found that 5 of the human centrosomal mRNAs had orthologs in *Drosophila*, and 4 of these localized to centrosomes in S2R+ cells. Moreover, the fifth *Drosophila* mRNA, Plp, was reported to localize to centrosomes in *Drosophila* embryos (Table 1; Lécuyer et al., 2007; Wilk et al., 2016). These data show a striking and unprecedented degree of evolutionary conservation in RNA localization, where the same family of mRNAs is localized to the same subcellular site, from *Drosophila* to humans. This likely underlies conserved features in mRNA localization mechanism and/or function.

### A cell-cycle dependent translational program operates at centrosomes

Our data reveal that centrosomal mRNA localization varies with phases of the cell cycle. Two mRNAs specifically localized in mitosis (ASPM and NUMA1), one in interphase but not mitosis (CCDC88C), and 5 in interphase and early mitosis (HMMR, BICD2, CEP350, PCNT, NIN). The phase where most mRNAs (7 out of 8) localize is thus prophase. Moreover, the two mitotic mRNAs localized with different kinetics: ASPM localized during all mitotic phases while NUMA1 only during prophase and prometaphase. Finally, HMMR was the only transcript that localized at the cytokinetic bridge at the end of cell division, together with its protein. Interestingly, the *Drosophila* centrosomal mRNAs also localize in a cell cycle-dependent way, indicating that this feature is also conserved during evolution. This shows the variety, complexity and precision of centrosomal mRNA localization, as well as its potential role during the centrosome cycle. These data demonstrate the existence of a unique and conserved translational program at centrosomes, which is cell cycle regulated.

It is interesting to speculate why these eight proteins and not others are locally translated. Since they function in centrosome/spindle maturation and that this occurs over short time periods, having optimal amounts at centrosomes at the right time point of the cell cycle is crucial. Interestingly, most of these proteins have relatively large sizes (more than 2000 aa, with the exception of HMMR and BICD2), and it would thus take some time to synthesize them. A local translational regulation may thus provide an efficient and rapid method for targeting them to the centrosome when needed. This might be particularly important in prophase where most mRNAs localize, because it is a relatively short phase and also the site of important changes in centrosome composition and function.

Another possibility is that mRNA accumulation at centrosomes plays a structural role. An emerging model is that phase separation helps the formation of centrosomes (Woodruff et al., 2018). Since RNAs are often critical components of phase separated condensates (Navarro et al., 2019), their accumulation at centrosomes could contribute to their formation. Finally, a likely and not exclusive possibility is that these 8 proteins need to be assembled co-translationally with their partners at the centrosome. Co-translational folding occurs for many proteins and this can be facilitated by the presence of a protein’s partner. This may further provide an elegant mechanism for RNA localization (see below).

### Centrosomal mRNA localization occurs by active transport of polysomes and requires the nascent protein

For all the eight centrosomal transcripts studied here, premature ribosome termination delocalized the mRNAs while freezing the ribosome and the nascent protein chain on the mRNA had no effect. In the case of a GFP-tagged ASPM mRNA, we further observed that preventing translation of the nascent ASPM protein via a stop codon abolished centrosomal RNA localization, indicating that translation is required in *cis*. Most importantly, polysomes coding for ASPM and NUMA1 were actively transported to centrosomes at rates compatible with motor-driven transport (0.5-1 μm/s). Together, this shows that centrosomal mRNA localization relies on an active mechanism driven by the nascent peptide. While RNA localization is often conceptualized as an RNA-driven process that transports silenced mRNAs, our data contradict this dogma and show that for centrosomal mRNAs, polysome transport mediated by the nascent protein is the rule.

These observations suggest that the nascent polypeptide contains a localization signal that would drive the polysome toward centrosomes. Interestingly, we found that ASPM polysomes are actively transported via two mechanisms. The first is motorized transport whereby a polysome slides on MTs. The second involves the pulling of an entire MT with an ASPM polysome attached to it. The N-terminal part of ASPM contains two calponin-homology domains that can bind MTs. It is possible that once these domains are translated, they bind MTs and cause the entire ASPM polysome to attach. In agreement with this possibility, local translation of ASPM at MTs can also be seen during interphase. Interestingly, all eight localized mRNAs encode proteins that either bind MTs directly (ASPM, NUMA1, HMMR, CCDC88C, CEP350) or contribute to MT anchoring (NIN, PCNT, BICD2). Similarly, many of these proteins directly or indirectly bind dynein (BICD2, NIN, CCDC88C, PCNT, NUMA1, HMMR; Redwine et al., 2017). These properties could thus be part of the transport mechanism.

The paradigm for translation-dependent RNA localization is that of secreted proteins. In this case, translation initially leads to the synthesis of the signal peptide, which is recognized by SRP. This halts the ribosome until the entire complex docks on the SRP receptor on the ER, where translation resumes. It is thus tempting to envision a scenario where ribosomes translating centrosomal mRNAs enter a pause and only resume translation after reaching the centrosome. It has been shown that unfolded domains can halt ribosomes (Liu et al., 2013; Shalgi et al., 2013). Moreover, in a recent case of co-translational assembly, the ribosome enters a pause at a specific location, which is relieved upon interaction with the partner of the nascent protein (Panasenko et al., 2019). If indeed the proteins encoded by the centrosomal mRNAs need to be co-translationally assembled, it is possible that the domain responsible for this would remain unfolded before the polysome reaches the centrosome. It could thus halt the ribosomes while the nascent polypeptide located upstream of this unfolded domain could connect the polysome to transport systems and drive it to the centrosome. This could be an elegant and general mechanism that ensures RNA localization and local translation, as it could work at any place in the cell. This could explain why co-translational mRNA targeting appears to be a widespread mechanism in cell lines (Chouaib et al., 2019).

## Acknowledgements

We thank A. Akhmanova for the ASPM cDNA, B. Delaval for the HeLa centrin1-GFP cell line, and the MGX and MRI facilities for their technical support. We thank Florian Müller and Thomas Walter for critical readings of the manuscript and scientific discussions, and Philippe Fort and Andreas Merdes for scientific inputs. AS was supported by fellowships from MESRI and FRM. This project was supported by France BioImaging (ANR-10-INBS-04), by the Agence Nationale de la Recherche (ANR-11-BSV8-018-02 and ANR-14-CE10-0018-01), the Institut Pasteur, the Ligue Nationale Contre le Cancer and the Labex EpiGenMed, from the framework “Investissements d’avenir”.

## Author contribution statement

The HT-smFISH methodology was conceived by EB and developed by AMT, CL, EC, EB, FL, MP, TG and VG. The idea of screening the centrosomal proteome was conceived by EB, the gene/probe lists were generated by CL, EB and MP, the screen was conducted by EC and FL, and image data interpretation/curation was performed by EC, AS and EB. OSK and HLH conceived and conducted investigations of translation factor localization. All other investigations were conceived and conducted by AS, with help from MCR and EB for the GFP-stop-ASPM CDS experiment. AS, AMT, EC, EB, FL, HLH, KZ, MCR, MP, OSK, RC, VG, and XP analyzed the data. AS prepared the Figures for data visualization. MP wrote the HT-smFISH method section, AS and EB wrote the abstract and discussion, and AS wrote the rest of the manuscript, which was reviewed and edited by AS, EB, HLH, KZ, MP, OSK and XP.

## Materials and Methods

### Cell lines, culture conditions and treatments

HeLa-Kyoto cells and HeLa cells expressing or not Centrin1-GFP (a gift from Dr. B. Delaval) were grown in Dulbecco’s modified Eagle’s Medium (DMEM, Gibco) supplemented with 10% fetal bovine serum (FBS, Sigma-Aldrich), and 100 U/mL penicillin/streptomycin (Sigma-Aldrich). The collection of HeLa-Kyoto cell lines stably transfected with the GFP-tagged BACs was described previously (Poser et al., 2008, Maliga et al., 2013), and these cells were grown in the same medium in addition to 400 μg/ml G418 (Gibco). All human cells were grown at 37°C with 5% CO_2_. S2R+ cells were grown in Schneider’s Drosophila medium (Gibco) supplemented with 10% fetal bovine serum (FBS, Sigma-Aldrich), and 100 U/mL penicillin/streptomycin (Sigma-Aldrich) at 25°C. Drugs were used at the following final concentrations: 100 μg/ml for puromycin, 200 μg/ml for cycloheximide, and 5 μg/ml for nocodazole. Treatment of cells with translation inhibitors was for 20 minutes (reduced to 5 min for mitotic cells when indicated). Treatment of cells with nocodazole was for 10 minutes. Transfection of the GFP-ASPM CDS constructs was done using JetPrime (Polyplus) and 2 μgs of DNA were transfected overnight in a 6 well plate containing 2ml of medium.

### Insertion of the MS2 cassette by CRISPR/Cas9

The recombination cassettes contained 500 bases of homology arms flanking a 3xHA tag, a stop codon and 24 MS2 repeats. A start codon was placed after the MS2 repeats followed by an IRES, a neomycin resistance gene, and a stop codon. The IRES-Neo^r^ segment was flanked by two LoxP sites having the same orientation. HeLa Kyoto cells were transfected using JetPrime (Polyplus) and a cocktail of four plasmids, including the recombination cassette and constructs expressing Cas9-nickase and two guide RNAs with an optimized scaffold (Pichon et al. 2016). Insertion was targeted at the stop codon of the ASPM and NUMA1 genes. Cells were selected on 400 μg/ml G418 neomycin for a few weeks. Individual clones were then picked and analyzed by PCR genotyping, fluorescent microscopy and smFISH/smiFISH with probes against both the endogenous ASPM or NUMA1 mRNA and MS2 sequences. Stable MCP-GFP-NLS expression was then set up via retroviral infection. The sequences targeted by the guide RNAs were (PAM sequences are underlined): TCTCTTCTCAAAACCCAATC<u>TGG</u> for ASPM guide 1, and GCAAGCTATTCAAATGGTGA<u>TGG</u> for ASPM guide 2; GAGGTCAGCATCGGGGACAC<u>AGG</u> for NUMA1 guide 1, and AGTGCCTTCTCTCAGCTCCC<u>AGG</u> for NUMA1 guide 2.

### Insertion of SunTag cassette by CRISPR/Cas9

The recombination cassettes contained 500 bases of homology arms flanking a puromycin resistance gene translated from the endogenous ATG sequence, followed by a P2A sequence, 32 SunTag repeats, and a P2A-T2A-FLAG sequence in the case of ASPM or a P2A-FLAG sequence in the case of NUMA1, fused to the protein of interest. Hela Kyoto cells stably expressing the scFv-GFP were transfected using JetPrime (Polyplus) and a cocktail of three plasmids, including the recombination cassette, and constructs expressing Cas9-HF1 and guide RNAs with an optimized scaffold. Cells were selected on 0.25 μg/ml puromycin for a few weeks. Individual clones were then picked and analyzed by PCR genotyping, fluorescent microscopy and smiFISH with probes against the SunTag or endogenous mRNA sequence. The sequences targeted by the guide RNAs were (PAM sequences are underlined): AAGTGAGCCCGACCGAGCGG<u>AGG</u> for ASPM and GACAGTCACTCCAATGCGCC<u>TGG</u> for NUMA1.

### Genotyping

PCR was done using a Platinum Taq DNA Polymerase (Invitrogen) on genomic DNA prepared with GenElute Mammalian Genomic DNA miniprep (Sigma-Aldrich). The sequences of oligonucleotides are given below.

For genotyping ASPM MS2×24 clones: 5’-TCAGAGGGTATGGAGGGGAA-3’ (ASPM gene end WT forward) with 5’-GACATCTGTGGCCCTGAAAC-3’ (ASPM gene end WT reverse) for the WT ASPM allele; and 5’-TCAGAGGGTATGGAGGGGAA-3’ (ASPM gene end WT forward) with 5’-GCCCTCACATTGCCAAAAGA-3’ (IRES reverse) for the edited ASPM allele.

For genotyping NUMA1 MS2×24 clones: 5’-ACCAAGGACTAAAGGGAGCC-3’ (NUMA1 gene end WT forward) with 5’-CAACCCCACTCCTGAGACAT-3’ (NUMA1 gene end WT reverse) for the WT NUMA1 allele; and 5’-ACCAAGGACTAAAGGGAGCC-3’ (NUMA1 gene end WT forward) with 5’-GCCCTCACATTGCCAAAAGA-3’ (IRES reverse) for the edited NUMA1 allele.

For genotyping SunTagx32 ASPM clones: 5’-TGTTCCTGGAAACCGCAATG-3’ (ASPM gene start WT forward) with 5’-GTTTATGTGTTGTCCCCGCC-3’ (ASPM gene start WT reverse) for the WT ASPM gene; and, 5’-AAAAGGGTAGCGGATCAGGA-3’ (SunTagx32 forward), with 5’-CATGTGTATGCGTCAAGGGC-3’ (ASPM gene start reverse) for the edited allele.

For genotyping SunTagx32 NUMA1 clones: 5’-TCATTGTGCCCCTGGAGATT-3’ (NUMA1 gene start WT forward) with 5’-CAGAGAGACCAGTGCTGTGA-3’ (NUMA1 gene start WT reverse) for the WT NUMA1 gene; and, 5’-ACCGGTGACTACAAAGACGA-3’ (FLAG forward), with 5’-GCTGTGATTCTATGCTGGGC-3’ (NUMA1 gene start reverse) for the edited allele.

### Single molecule fluorescent *in situ* hybridization

Cells grown on glass coverslips or 96-well glass bottom plates (SensoPlates, Greiner) were fixed for 20 min at RT with 4% paraformaldehyde (Electron Microscopy Sciences) diluted in PBS (Invitrogen), and permeabilized with 70% ethanol overnight at 4°C.

For smFISH performed on BAC cells, we used a set of 44 amino-modified oligonucleotide probes against the GFP-IRES-Neo sequence present in the BAC construction (sequences given in Table S2). Each oligonucleotide probe contained 4 primary amines that were conjugated to Cy3 using the Mono-Reactive Dye Pack (PA23001, GE Healthcare Life Sciences). To this end, the oligos were precipitated with ethanol and resuspended in water. For labelling, 4 μg of each probe was incubated with 6 μl of Cy3 (1/5 of a vial resuspended in 30 μl of DMSO), and 14 μl of carbonate buffer 0.1 M pH 8.8, overnight at RT and in the dark, after extensive vortexing. The next day, 10 μg of yeast tRNAs (Sigma-Aldrich) were added and the probes were precipitated several times with ethanol until the supernatant lost its pink color. For hybridization, fixed cells were washed with PBS and hybridization buffer (15% formamide from Sigma-Aldrich in 1X SSC), and then incubated overnight at 37°C in the hybridization buffer also containing 130 ng of the probe set for 100 μl of final volume, 0.34 mg/ml tRNA, 2 mM VRC (Sigma-Aldrich), 0.2 mg/ml RNAse-free BSA (Roche Diagnostics), and 10% Dextran sulfate (MP Biomedicals). The next day, the samples were washed twice for 30 minutes in the hybridization buffer at 37°C, and rinsed in PBS. Coverslips were then mounted using Vectashield containing DAPI (Vector laboratories, Inc.). For smFISH against the MS2 tag, 25 ng of an oligonucleotide labeled by two Cy3 molecules at the first and last thymidine (sequence in Table S2) was used per 100μl of hybridization mix.

For smiFISH using DNA probes (Tsanov et al., 2016), 24-48 unlabeled primary probes were used (sequences given in Table S2). In addition to hybridizing to their targets, these probes contained a FLAP sequence that was pre-hybridized to a secondary fluorescent oligonucleotide. To this end, 40 pmoles of primary probes were pre-hybridized to 50 pmoles of secondary probe in 10 μl of 100 mM NaCl, 50 mM Tris-HCl, 10 mM MgCl2, pH 7.9. Pre-hybridization was performed on a thermocycler with the following program: 85°C for 3 min, 65°C for 3 min, and 25°C for 5 min. The final hybridization mixture contained the probe duplexes (2 μl per 100 μl of final volume), with 1X SSC, 0.34 mg/ml tRNA, 15% Formamide, 2 mM VRC, 0.2 mg/ml RNAse-free BSA, 10% Dextran sulfate. Slides were then processed as above.

For smiFISH using RNA probes in the high-throughput smFISH screen, a pool of DNA oligonucleotides (GenScript) was used to generate the primary probes. The oligonucleotide design was based on the Oligostan script (Tsanov et al., 2016). A first series of PCR was performed using the oligopool as template and each of the gene-specific barcoding primers. A second series of PCR was achieved using the following primers: FLAP Y sequence with the addition of the T7 RNA polymerase promoter sequence at its 5’ end (5’ TAATACGACTCACTATAGGGTTACACTCGGACCTCGTCGACATGC ATT-3’), and reverse complement sequence of FLAP X (5’-CACTGAGTCCAGCTCGA AACTTAGGAGG-3’). All PCR reactions were carried out with Phusion DNA Polymerase (Thermo Fisher Scientific), in 96-well plates with a Freedom EVO 200 (Tecan) robotic platform. PCR products were checked by capillary electrophoresis on a Caliper LabChip GX analyzer (PerkinElmer). The products of the second PCR were purified with a NucleoSpin 96 PCR Clean-up kit (Macherey-Nagel), lyophilized, and resuspended in DNase/RNase-free distilled water (Invitrogen). *In vitro* transcription was subsequently performed with T7 RNA Polymerase and the obtained primary probes were analyzed by capillary electrophoresis using a Fragment Analyzer instrument (Advanced Analytical). 50 ng of primary probes (total amount of the pool of probes) and 25 ng of each of the secondary probes (TYE 563 labeled LNA oligonucleotides targeting FLAP X and FLAP Y, Qiagen) were pre-hybridized in 100 μL of the following buffer : 1X SSC, 7.5 M urea (Sigma-Aldrich), 0.34 mg/mL tRNA, 10% Dextran sulfate. Pre-hybridization was performed on a thermocycler with the following program: 90°C for 3 min, 48°C for 15 min. Plates with fixed cells were washed with PBS and hybridization buffer (1X SSC, 7.5 M urea). Hybridization was then carried out overnight at 48°C. The next day, plates were washed six times for 10 minutes in 1xSSC 7.5M urea at 48°C. Cells were rinsed with PBS at RT, stained with 1 μg/mL Dapi diluted in PBS, and mounted in 90% glycerol (VWR), 1 mg/mL p-Phenylenediamine (Sigma-Aldrich), PBS pH 8.

### Immunofluorescence

S2R+ cells were seeded and fixed as for smFISH. Cells were permeabilized with 0.1% Triton-X100 in PBS for 10 minutes at room temperature and washed twice with PBS. For centrosome labelling, coverslips were incubated for one hour at room temperature with a monoclonal anti-γ-tubulin antibody produced in mouse (Sigma-Aldrich, T5326), diluted 1/1000 in PBS. Coverslips were washed twice with PBS, 5 minutes each time, and incubated with either an FITC (Jackson ImmunoResearch 115-095-062) or Cy5 (Jackson ImmunoResearch 115-176-003) labeled anti-mouse secondary antibody diluted 1/100 in PBS. After 1 hour of incubation at RT, coverslips were washed twice with PBS, 5 minutes each. Coverslips were mounted using Vectashield containing DAPI (Vector laboratories, Inc.).

### Imaging of fixed cells

Microscopy slides were imaged on a Zeiss Axioimager Z1 wide-field microscope equipped with a motorized stage, a camera scMOS ZYLA 4.2 MP, using a 63x or 100x objective (Plan Apochromat; 1.4 NA; oil). Images were taken as Z-stacks with one plane every 0.3 μm. The microscope was controlled by MetaMorph and figures were constructed using ImageJ, Adobe Photoshop and Illustrator.

96-well plates were imaged on an Opera Phenix High-Content Screening System (PerkinElmer), with a 63x water-immersion objective (NA 1.15). Three-dimensional images were acquired, consisting of 35 slices with a spacing of 0.3 μm.

### Image analysis and quantifications

Mitotic phases were identified based on visual inspection of DNA condensation and cell shape. Early prophase was defined by its low DNA compaction, which increased in late prophase. Early prometaphase was marked by the rupture of the nuclear envelope, while late prometaphase additionally displayed cell rounding. For late mitosis, we subdivided cells into early telophase (without cytokinesis), and late telophase (with cytokinesis marked by the accumulation of HMMR-GFP at cytokinetic bridges). Centrosomal localization was assessed by visual inspection of individual cells.

### Imaging of live cells

Live imaging was done using a spinning disk confocal microsope (Nikon Ti with a Yokogawa CSU-X1 head) operated by the Andor iQ3 software. Acquisitions were performed using a 100X objective (CF1 PlanApo λ 1.45 NA oil), and an EMCCD iXon897 camera (Andor).

For fast imaging, we imaged at a rate of at least 1 stack/s for 1-3 mins, using stacks with a Z-spacing of 0.4-0.6 μm. This spacing allowed accurate point spread function determination without excessive oversampling. For slow imaging, we collected stacks of around 19 planes with a Z-spacing of 0.6 μm and at a frame rate of one stack every 5 mins for 62 hours. The power of illuminating light and the exposure time were set to the lowest values that still allowed visualization of the signal. This minimized bleaching, toxicity and maximized the number of frames that were collected. Samples were sequentially excited at 488 and 640 nm in the case of dual-color imaging.

For ASPM-MS2×24 and NUMA1-MS2×24, a time lapse of a Z-stack covering a 3D section of the cell was acquired in the 488 nm channel while a single Z-stack was acquired in the 640 nm channel to identify the mitotic phase. For mono-color SunTag ASPM and NUMA1 movies, Z-stacks traversing the entire cell were imaged. For dual color imaging of SunTag-ASPM and MTs, a Z-stack covering a 3D section of the cell was imaged, to maintain high frame rates and compensate for time required for the second color.

Cells were maintained in anti-bleaching live cell visualization medium (DMEM^gfp^; Evrogen), supplemented with 10% fetal bovine serum at 37°C in 5% CO_2_. SiR-DNA (Spirochrome) was kept at 100 nM throughout the experiments to label DNA. SiR-tubulin (Spirochrome) was kept at 100 nM throughout the experiments to label MTs.

### Single molecule dynamics analysis and single particle tracking

The dynamics of ASPM mRNAs, and both NUMA1 mRNAs and polysomes were assessed as follows: mRNAs anchored to the centrosomes as well as those undergoing directed motion were identified based on visual inspection. The speed of individual mRNA molecules displaying directed motions was measured across at least 3 frames and calculated using ImageJ.

Single particle tracking of ASPM polysomes was performed using the TrackMate plugin in ImageJ. The DoG detector was used. Blob diameter was set to 0.7-0.8 microns and the detection threshold was between 100-120. Median filtering and sub-pixel localization were additionally used. The simple LAP tracker option was used to construct tracks. Both linking and gap closing distances were assigned a maximum value of 1.5 microns. Three frame gaps were allowed when constructing tracks. Tracks were displayed color-coded according to displacement (red corresponds to highest values while blue corresponds to lowest). For SunTag-ASPM mono-color movies, the top 20 tracks with highest displacements were chosen. The velocity was calculated by measuring the displacement over at least 5 frames of directed motion. Directionality towards the centrosome was determined visually. For dual color movies of SunTag-ASPM and MTs, tracks shorter than 15 seconds were filtered out. Tracks were classified in one of three categories based on their localization (colocalizing to MT, colocalizing with the nuclear envelope), while a fourth category was made for particles showing directed motions.

Tracks were imported and analyzed in R. Instant 1D displacements between frames were calculated along the x and y axis, and the resulting histograms were fitted to a Gaussian function, for which variance is directly proportional to the diffusion coefficient *(D).* We also calculated a mean MSD as a function of time, by aligning all tracks at their start and averaging the resulting 2D displacements.

## Supplementary spreadsheets

**Table S1:** Summary of all screened mRNAs

**Table S2:** Sequences of all smFISH and smiFISH probes

## Movie legends

**Movie 1**: HeLa cells expressing the edited ASPM-MS2×24 and MCP-GFP-NLS were imaged every 0.7 seconds for around 70 seconds during metaphase. The GFP signal shown in black corresponds to ASPM-MS2×24 mRNAs. The red arrow follows an mRNA showing directed movement.

**Movie 2**: HeLa cells expressing the edited ASPM-MS2×24 mRNA and MCP-GFP-NLS were imaged every 0.625 seconds for around 120 seconds during prometaphase. The GFP signal shown in green corresponds to ASPM-MS2×24 mRNAs; the Cy5 signal shown in red corresponds to DNA and was imaged for a single frame.

**Movie 3**: HeLa cells expressing the edited SunTagx32-ASPM allele and scFv-sfGFP were imaged every 0.9 seconds for 180 seconds during prophase. The GFP signal shown in black corresponds to ASPM polysomes and mature protein.

**Movie 4**: HeLa cells expressing the edited SunTagx32-ASPM allele and scFv-sfGFP were imaged every 0.9 seconds for 180 seconds during prometaphase. The GFP signal shown in black corresponds to ASPM polysomes and mature protein.

**Movie 5**: HeLa cells expressing the edited SunTagx32-ASPM allele and scFv-sfGFP were imaged every 0.66 seconds for 130 seconds during interphase with labeled MTs. The GFP signal shown in green corresponds to ASPM polysomes and mature protein; the Cy5 signal shown in red corresponds to MTs.

**Movie 6**: HeLa cells treated with nocodazole, expressing the edited SunTagx32-ASPM allele and scFv-sfGFP were imaged every 0.53 seconds for 105 seconds during interphase with labeled MTs. The GFP signal shown in green corresponds to ASPM polysomes and mature protein; the Cy5 signal shown in red corresponds to MTs and centrosomes.

**Movie 7**: HeLa cells expressing the edited SunTagx32-ASPM allele and scFv-sfGFP were imaged every 0.66 seconds for 260 seconds during prophase with labeled MTs. The GFP signal shown in green corresponds to ASPM polysomes and mature protein; the Cy5 signal shown in red corresponds to MTs.

**Movie 8**: HeLa cells expressing the edited SunTagx32-ASPM allele and scFv-sfGFP were imaged every 0.66 seconds for 130 seconds during prometaphase with labeled MTs. The GFP signal shown in green corresponds to ASPM polysomes and mature protein; the Cy5 signal shown in red corresponds to MTs.

**Movie 9**: HeLa cells expressing the edited NUMA1-MS2×24 allele and MCP-GFP-NLS were imaged every 0.9 seconds for around 80 seconds during prometaphase. The GFP signal shown in black corresponds to NUMA1 mRNA. The red arrow follows an mRNA showing directed movement.

**Movie 10**: HeLa cells expressing the edited SunTagx32-NUMA1 allele and scFv-sfGFP were imaged every 0.74 seconds for around 180 seconds during prometaphase. The GFP signal shown in black corresponds to ASPM polysomes and mature protein.

