## Supplemental figures for "A conserved choreography of mRNAs at centrosomes reveals a localization mechanism involving active polysome transport"

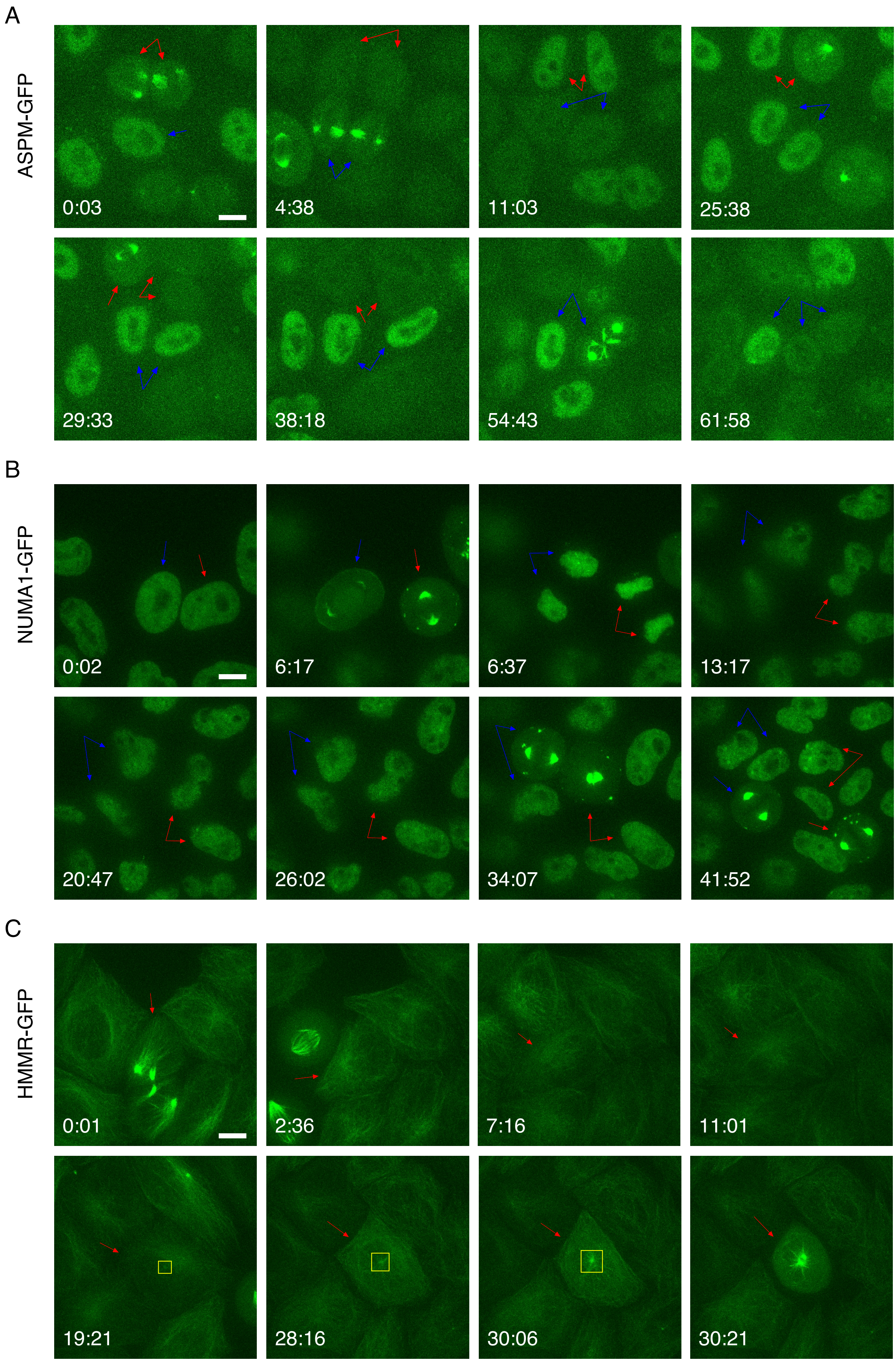


Figure S1

Safieddine et al.

**Figure S1 (related to Figure 2): Expression and localization of ASPM, NUMA1, and HMMR proteins across an entire cell cycle.**

(A) Images are snapshots of living HeLa cells expressing an ASPM-GFP BAC and imaged across an entire cell cycle. Signal is in green and corresponds to the ASPM-GFP protein. Scale bar is 10 microns and time is in hours: minutes. Red and blue arrows follow two dividing cells.

(B) Legend as in A, but for HeLa cells containing a NUMA1-GFP BAC.

(C) Legend as in A, but for HeLa cells containing a HMMR-GFP BAC.

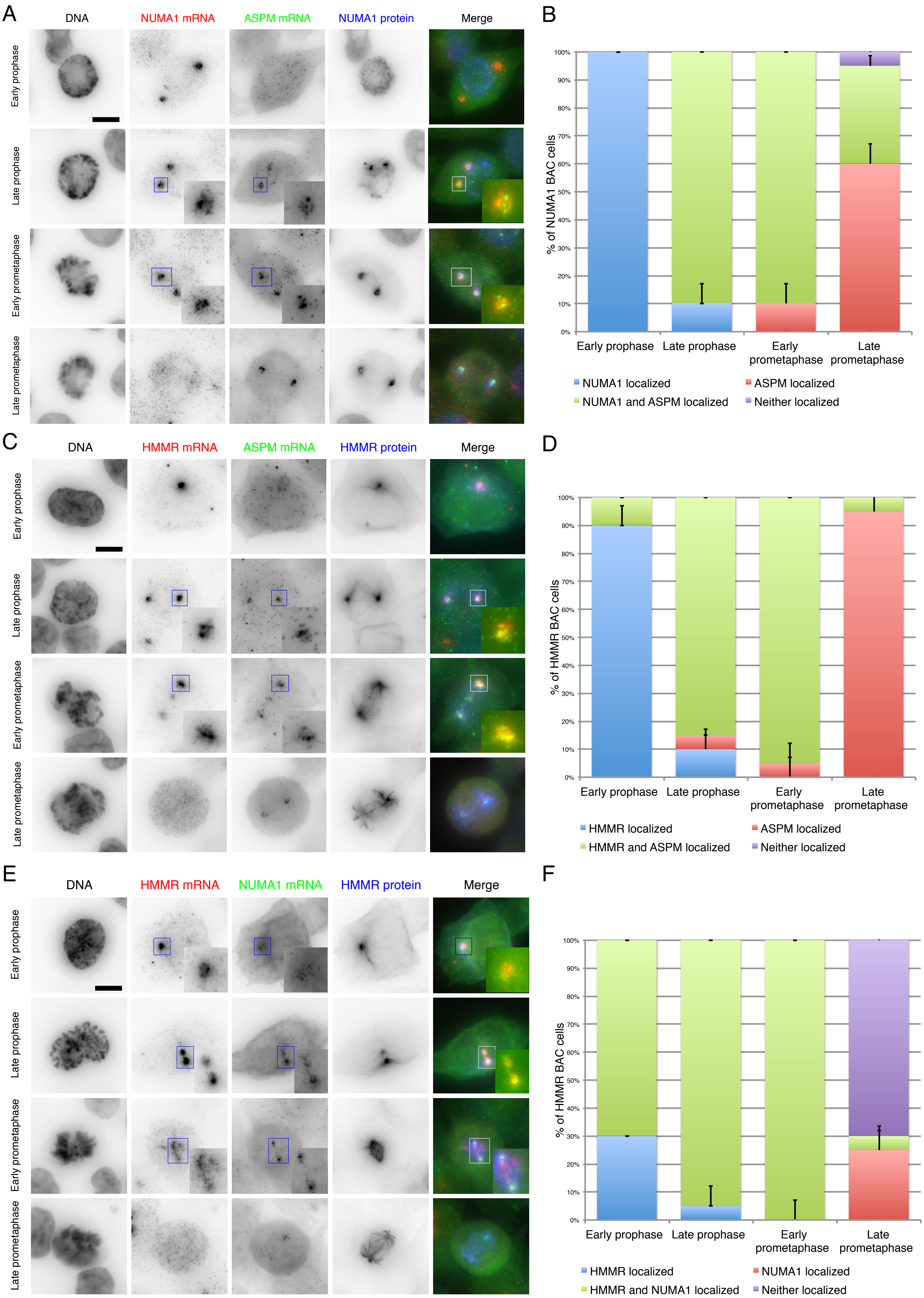


Figure S2

Safieddine et al.

**Figure S2 (related to Figure 2): ASPM, NUMA1, and HMMR mRNAs localize to distinct peri-centrosomal regions and at precise times during early cell division**.

(A) Images are micrographs of HeLa cells containing a NUMA1-GFP BAC and captured at different phases of early mitosis. Middle left and red: Cy3 fluorescent signals corresponding to NUMA1-GFP mRNAs labeled by smFISH; middle right and green: Cy5 signals corresponding to ASPM mRNA labeled by smiFISH. DNA stained with DAPI. Scale bar: 10 microns. Insets represent zooms of the boxed areas.

(B) Stacked histogram showing the percentage of cells having one, both, or neither transcript localized to centrosomes in each phase (n=10 cells per phase, repeated twice). Data represent mean and standard deviation.

(C and D) Legend as in A and B, but for HeLa cells containing an HMMR-GFP BAC with HMMR-GFP mRNA labeled in Cy3 shown in red, and ASPM mRNA labeled in Cy5 and shown in green.

(E and F) Legend as in A and B, but for HeLa cells containing an HMMR-GFP BAC with HMMR-GFP mRNA labeled in Cy3 shown in red, and NUMA1 mRNA labeled in Cy5 and shown in green.

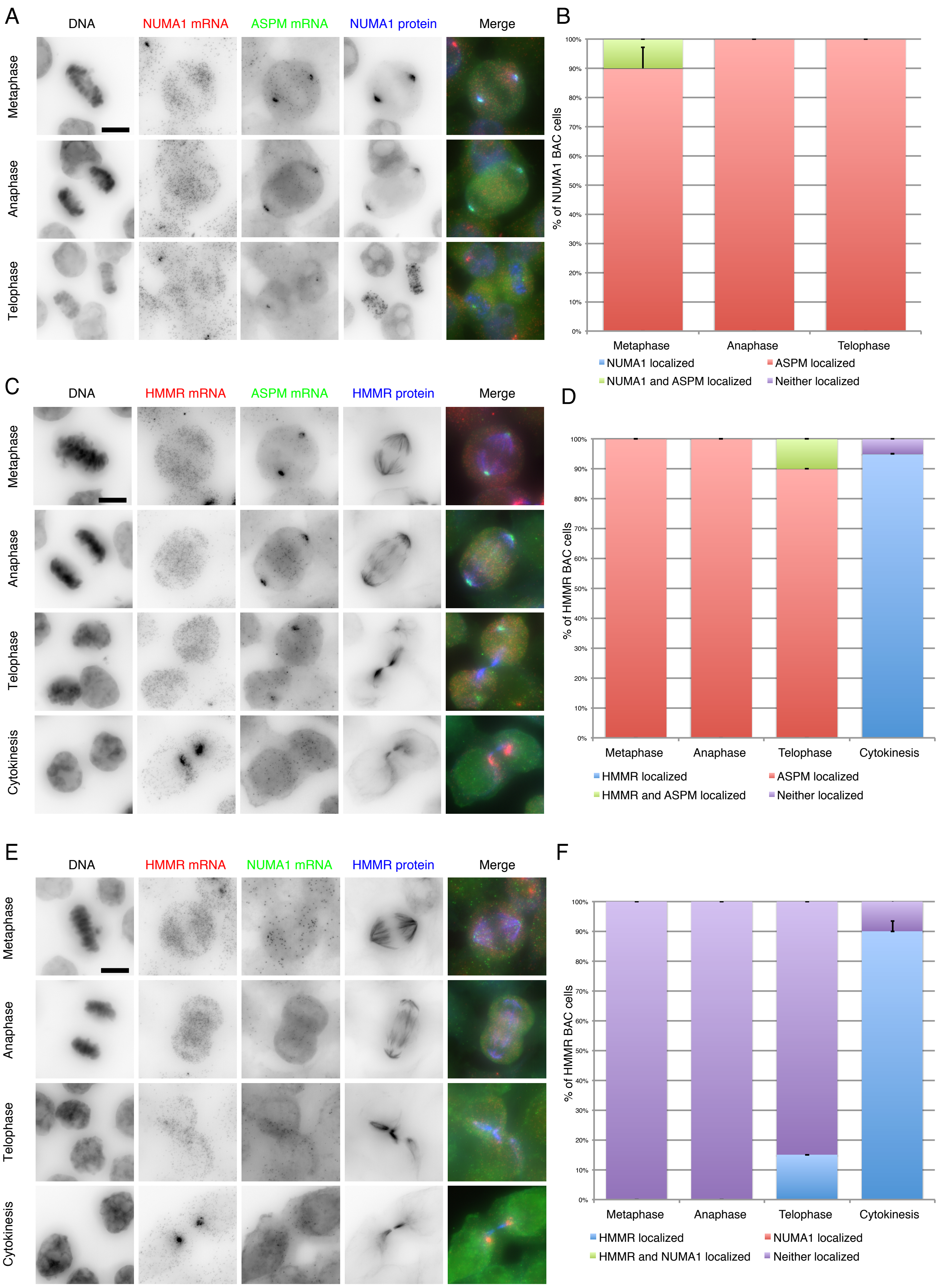


Figure S3

Safieddine et al.

**Figure S3 (related to Figure 2): ASPM, NUMA1, and HMMR mRNAs differentially localize to centrosomes during late cell division.**

(A) Images are micrographs of HeLa cells containing a NUMA1-GFP BAC and captured at different phases of late mitosis. Middle left and red: Cy3 fluorescent signals corresponding to NUMA1 mRNAs labeled by smFISH; middle right and green: Cy5 signals corresponding to ASPM mRNA labeled by smiFISH. DNA stained with DAPI. Scale bar: 10 microns.

(B) Stacked histogram showing the percentage of cells having one, both, or neither transcript localized to centrosomes in each phase (n=10 cells per phase, repeated twice). Data represent the mean and standard deviation.

(C and D) Legend as in A and B, but for HeLa cells containing an HMMR-GFP BAC with HMMR mRNA labeled in Cy3 shown in red, and ASPM mRNA labeled in Cy5 and shown in green.

(E and F) Legend as in A and B, but for HeLa cells containing an HMMR-GFP BAC with HMMR mRNA labeled in Cy3 shown in red, and NUMA1 mRNA labeled in Cy5 and shown in green.

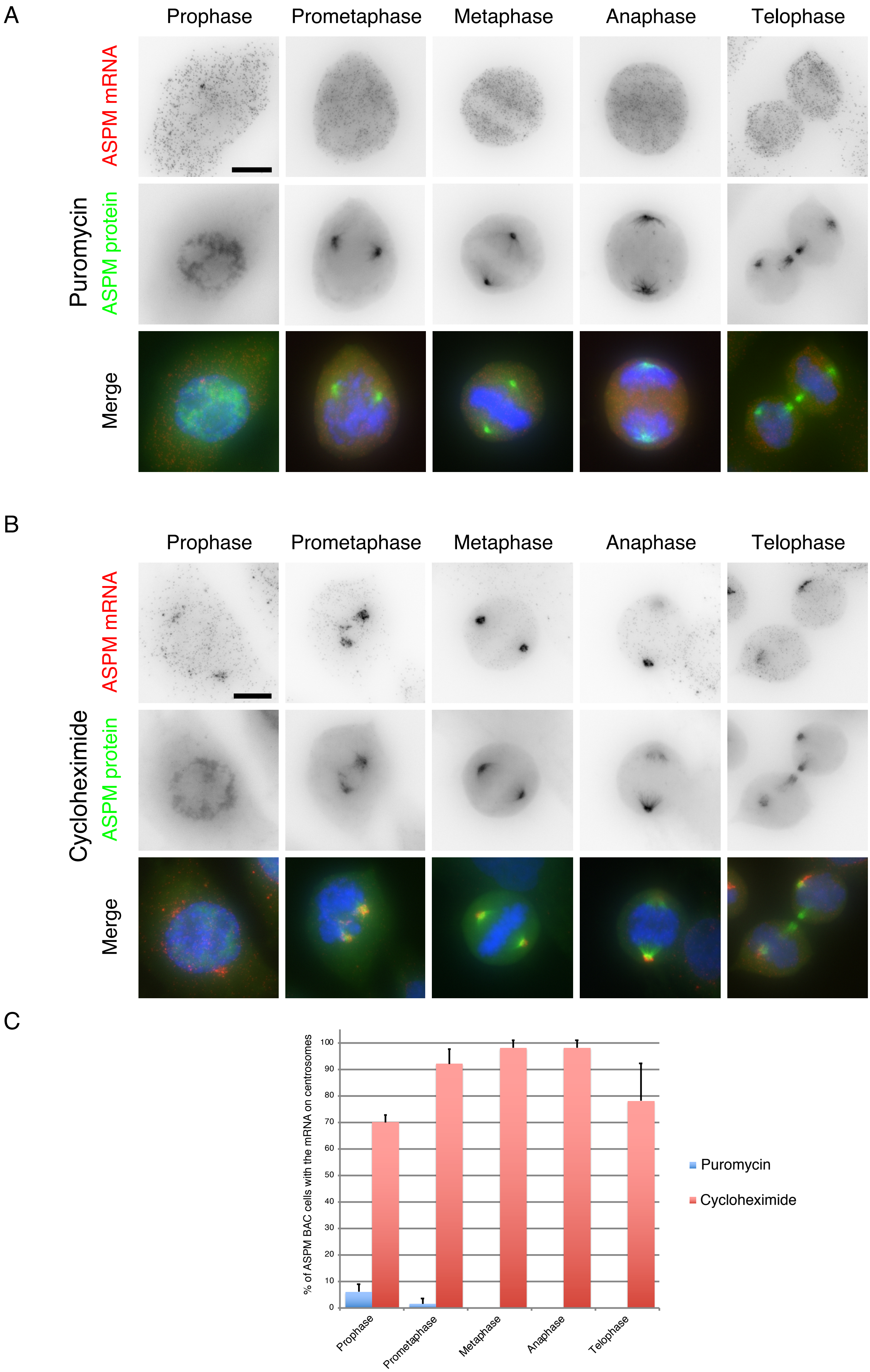


Figure S4

Safieddine et al.

**Figure S4 (related to Figure 3): Translation initiation is required for the localization of ASPM mRNAs during all phases of mitosis.**

(A) Micrographs of HeLa cells expressing an ASPM-GFP BAC, treated with puromycin and imaged during mitosis. Up and red: Cy3 fluorescent signals corresponding to ASPM mRNAs labeled by smFISH with probes against the GFP RNA sequence; middle and green: GFP signals corresponding to the ASPM protein. Blue: DNA stained with DAPI. Scale bar: 10 microns.

(B) Legend same as in A, but for cells treated with cycloheximide.

(C) Histogram depicting the percentage of cells showing a centrosomal localization of ASPM-GFP mRNAs after the indicated treatment (n=25 cells per phase, repeated twice). Data represent the mean and standard deviation.

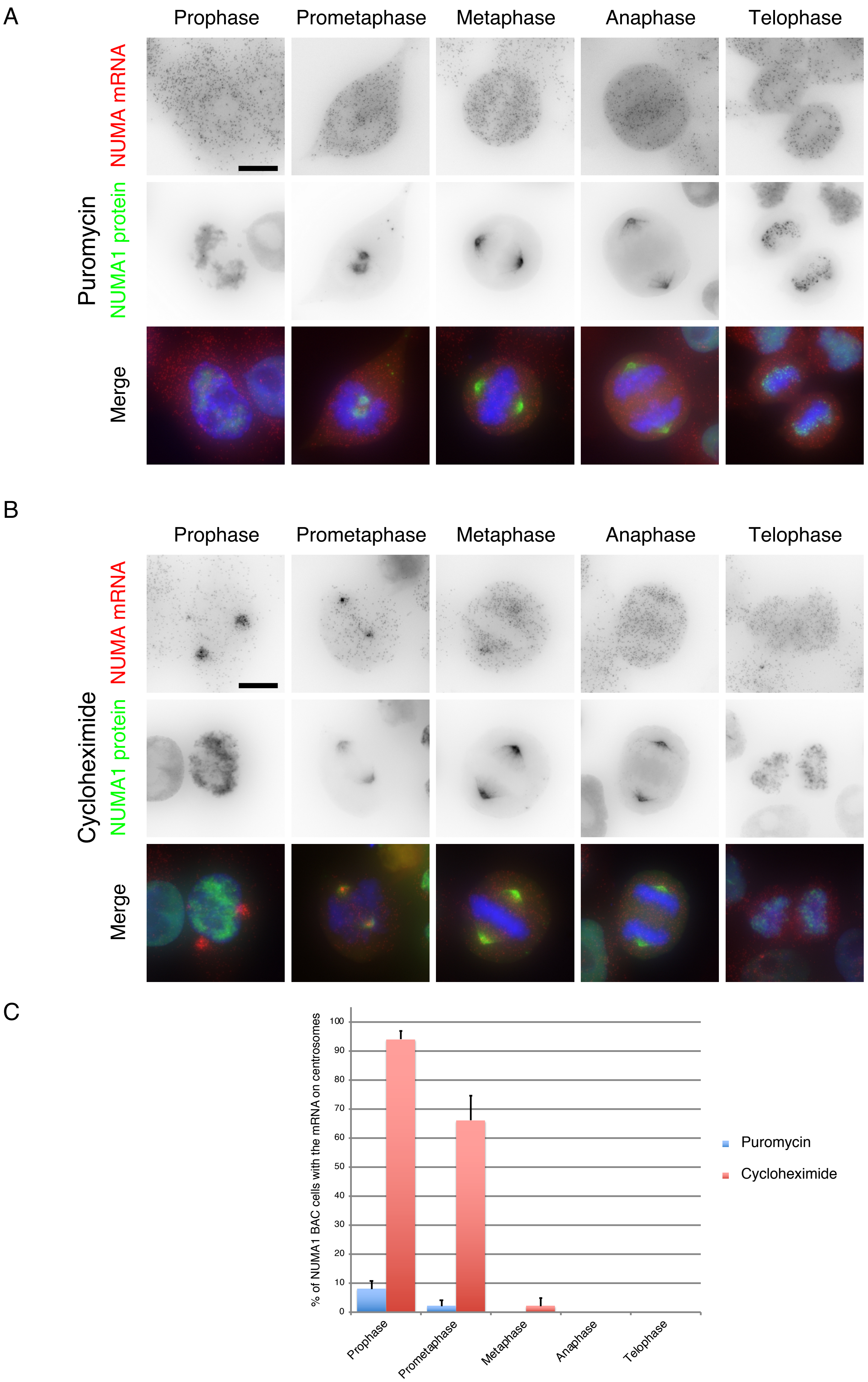
A
Prophase
Prometaphase
Metaphase
Anaphase
Telophase
Puromycin
NUMA mRNA
NUMA1 protein
Merge
B
Prophase
Prometaphase
Metaphase
Anaphase
Telophase
Cycloheximide
NUMA mRNA
NUMA1 protein
Merge
C
% of NUMA1 BAC cells with the mRNA on centrosomes
100
90
80
70
60
50
40
30
20
10
0
Prophase
Prometaphase
Metaphase
Anaphase
Telophase
Puromycin
Cycloheximide

Figure S5

**Figure S5 (related to figure 3): Translation initiation is required for the localization of NUMA1 mRNAs during all phases of mitosis.**

(A) Micrographs of HeLa cells expressing a NUMA1-GFP BAC, treated with puromycin and imaged during all phases of mitosis. Up and red: Cy3 fluorescent signals corresponding to NUMA1-GFP mRNA labeled by smFISH with probes against the GFP RNA sequence; middle and green: fluorescent signals corresponding to the NUMA1-GFP protein. Blue: DNA stained with DAPI. Scale bar: 10 microns.

(B) Legend same as in A, but for cells treated with cycloheximide.

(C) Histogram depicting the percentage of cells showing centrosomal localization of NUMA1-GFP mRNA after the indicated treatment (n=25 cells per phase, repeated twice). Data represent the mean and standard deviation.

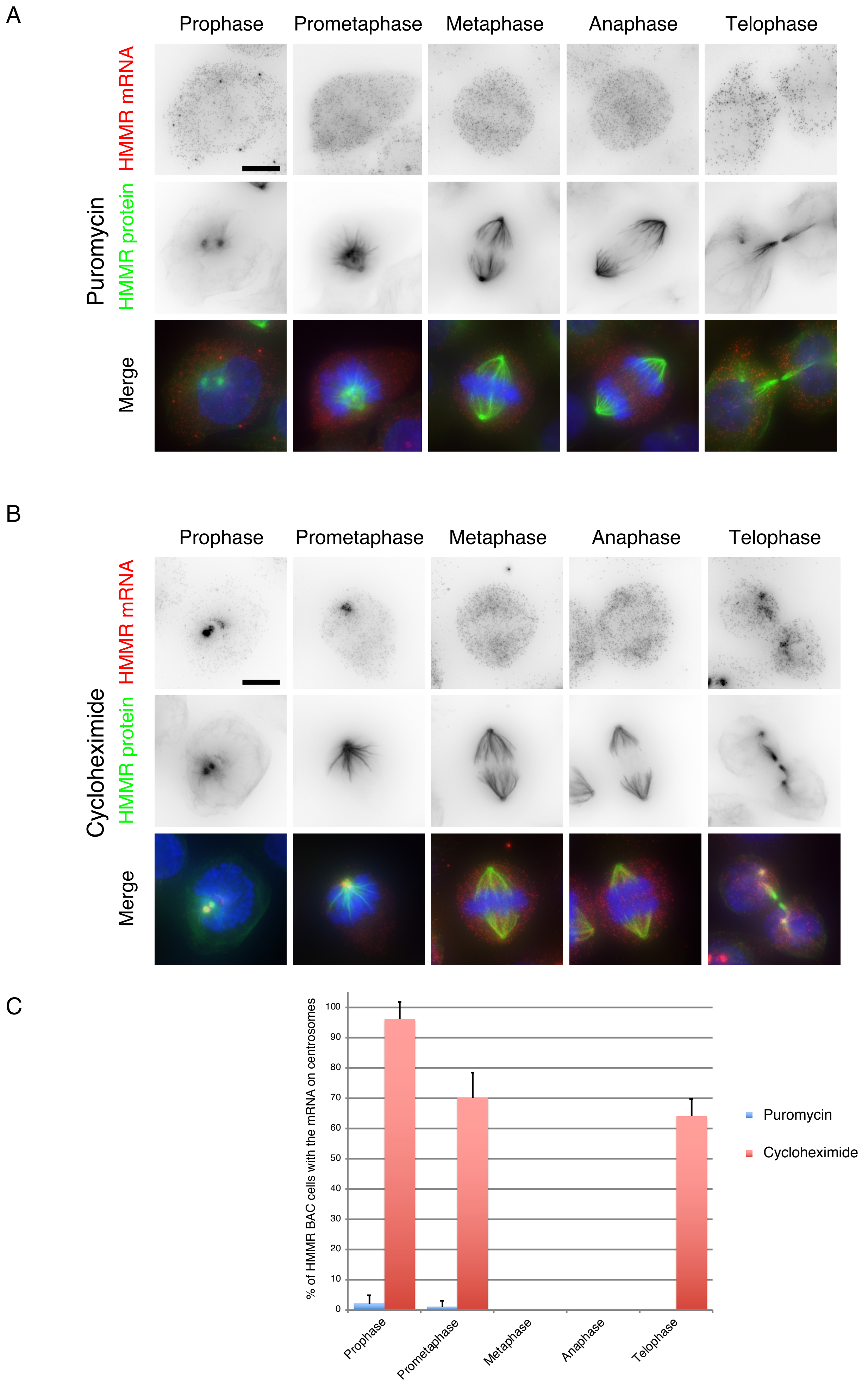


Figure S6

Safieddine et al.

**Figure S6 (related to Figure 3): Translation initiation is required for the localization of HMMR mRNAs during all phases of mitosis.**

(A) Micrographs of HeLa cells expressing a HMMR -GFP BAC, treated with puromycin and imaged during mitosis. Up and red: Cy3 fluorescent signals corresponding to HMMR-GFP mRNAs labeled by smFISH with probes against the GFP RNA sequence; middle and green: fluorescent signals corresponding to the HMMR-GFP protein. Blue: DNA stained with DAPI. Scale bar: 10 microns.

(B) Legend same as in A, but for cells treated with cycloheximide.

(C) Histogram depicting the percentage of cells showing centrosomal localization of HMMR-GFP mRNA after the indicated treatment (n=25 cells per phase, repeated twice). Data represent the mean and standard deviation.

A

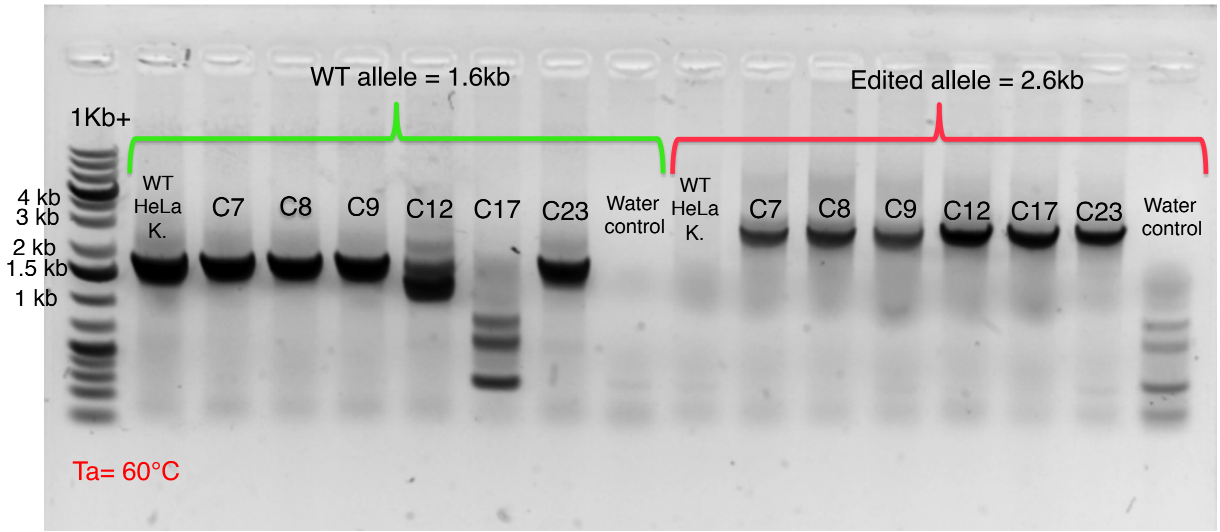
WT allele = 1.6kb
Edited allele = 2.6kb
1Kb+
4 kb
3 kb
2 kb
1.5 kb
1 kb
WT HeLa K.
C7
C8
C9
C12
C17
C23
Water control
WT HeLa K.
C7
C8
C9
C12
C17
C23
Water control
Ta= 60°C

B

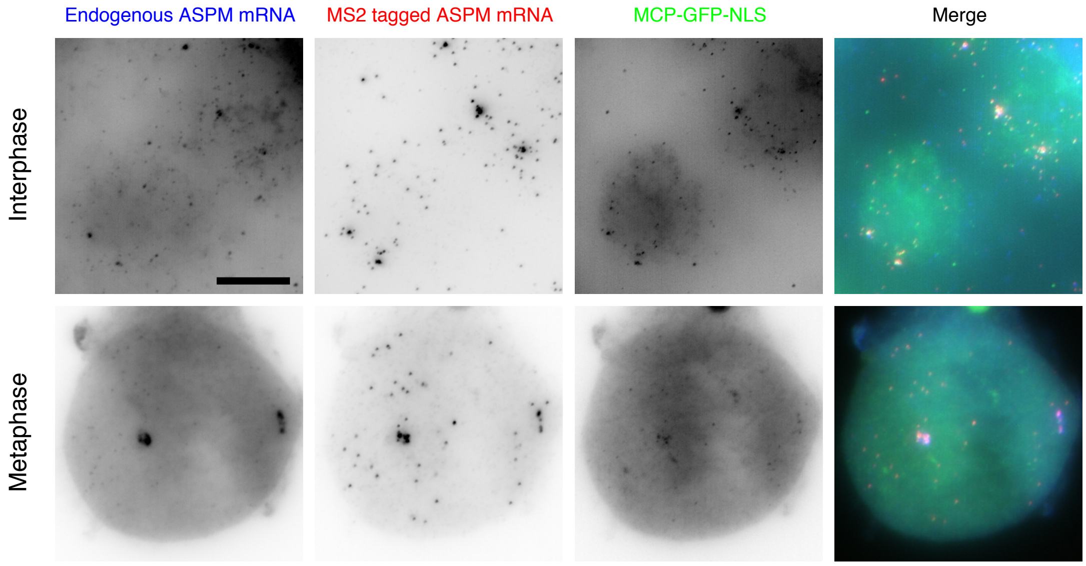
Endogenous ASPM mRNA
MS2 tagged ASPM mRNA
MCP-GFP-NLS
Merge
Interphase
Metaphase

Figure S7

**Figure S7 (related to Figure 5): Characterization of the ASPM-MS2x24 clone.**

(A) Image is a scan of a gel loaded with the product of a PCR performed on genomic DNA extracted from various ASPM-MS2x24 CRISPR clones. Wild-type and edited alleles are differentially amplified and give a product size of 1.6 and 2.6 kb, respectively. A ladder is placed on the left with the corresponding size markers. WT HeLa K.: PCR performed on the parental unedited cell line. Water control: PCR performed without any DNA. Ta: annealing temperature.

(B) Micrographs of ASPM-MS2x24 HeLa cells expressing MCP-GFP-NLS and imaged at interphase and mitosis. Left and blue: Cy5 fluorescent signals corresponding to tagged and untagged ASPM mRNAs labeled by smiFISH with probes against the endogenous mRNA; middle and red: Cy3 signals corresponding to tagged mRNAs labeled by smFISH with probes against the MS2 sequence; right and green: GFP signals corresponding to ASPM-MS2x24 mRNAs labeled by the MCP-GFP-NLS. Blue: DNA stained with DAPI. Scale bar: 10 microns.

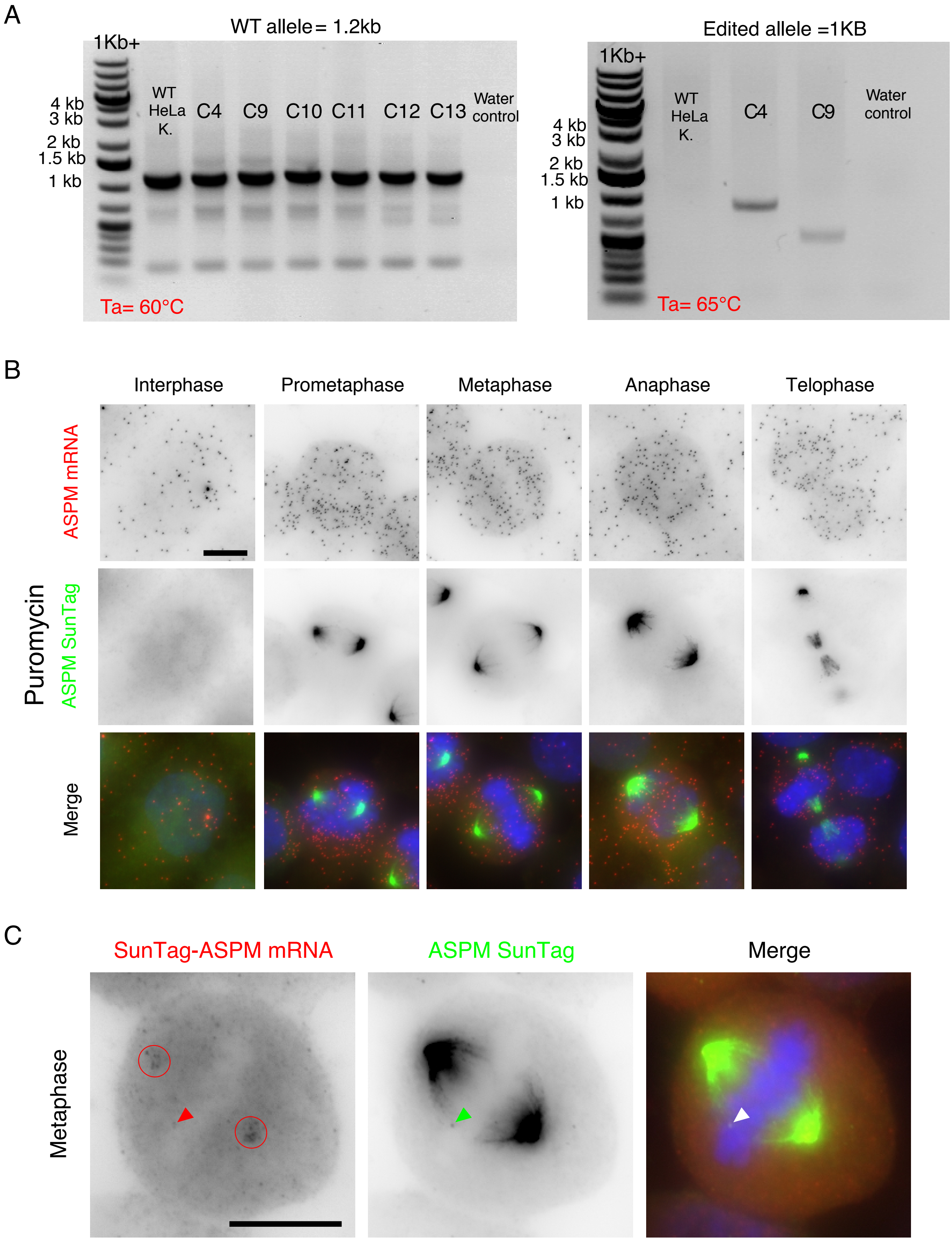
A
WT allele= 1.2kb
1Kb+
4 kb
3 kb
2 kb
1.5 kb
1 kb
WT HeLa K.
C4
C9
C10
C11
C12
C13
Water control
Ta= 60°C
Edited allele =1KB
1Kb+
4 kb
3 kb
2 kb
1.5 kb
1 kb
WT HeLa K.
C4
C9
Water control
Ta= 65°C
B
Interphase
Prometaphase
Metaphase
Anaphase
Telophase
ASPM mRNA
Puromycin
ASPM SunTag
Merge
C
SunTag-ASPM mRNA
ASPM SunTag
Merge
Metaphase

Figure S8

**Figure S8 (related to Figure 6): Characterization of the SunTagx32-ASPM clone**

(A) Image is a scan of a gel loaded with the product of a PCR performed on genomic DNA extracted from various SunTagx32-ASPM clones. Wild-type and edited alleles are differentially amplified and give a product size of 1.2 and 1 kb, respectively. A ladder is placed on the left with the corresponding size markers. WT HeLa K.: PCR performed on the parental unedited cell line. Water control: PCR performed without any DNA. Ta: annealing temperature

(B) Micrographs of SunTagx32-ASPM HeLa cells expressing the scFv-sfGFP, treated with puromycin and imaged during interphase and mitosis. Upper and red: Cy3 fluorescent signals corresponding to tagged and untagged ASPM mRNAs labeled by smFISH; middle and green: GFP signals corresponding to the SunTagx32-ASPM mature protein. Blue: DNA stained with DAPI. Scale bar: 10 microns.

(C) Images are micrographs of a HeLa clone expressing endogenous SunTagx32-ASPM and scFv-sfGFP. Left and red: Cy3 fluorescent signals corresponding to ASPM mRNA tagged with 32 SunTag repeats revealed by smFISH against the SunTag and puromycin sequences; middle and green: GFP signals corresponding to the ASPM mature protein. Blue: DNA stained with DAPI. Scale bar: 10 microns. The red and green arrows indicate an mRNA and a polysome respectively. The red circles indicate clusters of tagged ASPM mRNA.

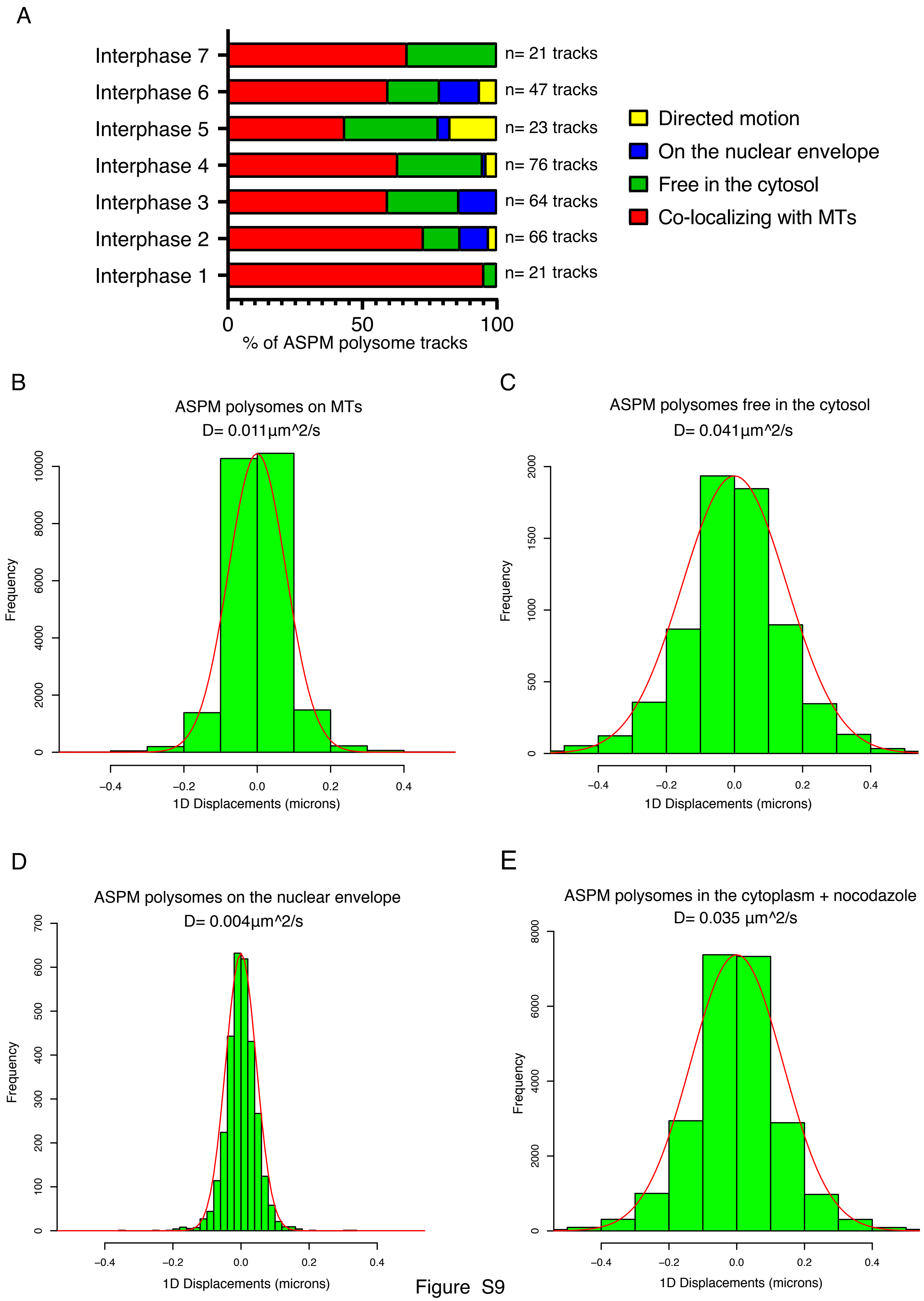


Figure S9

Safieddine et al.

**Figure S9 (related to Figure 7): 1D displacements of ASPM polysomes**

(A) Stacked bar graphs showing the proportion of the ASPM polysome tracks in the indicated categories.

(B) Histogram showing 1D displacements measured between two consecutive time frames of ASPM polysome, for tracks localizing on MTs. D is the average diffusion coefficient.

(C) Legend same as in B, but for ASPM polysome tracks free in the cytosol (not on MTs or the nuclear envelope)

(E) Legend same as in B, but for ASPM polysome tracks on the nuclear envelope.

(F) Legend same as in B, but for cells treated with nocodazole and ASPM polysomes not localizing at the nuclear envelope.

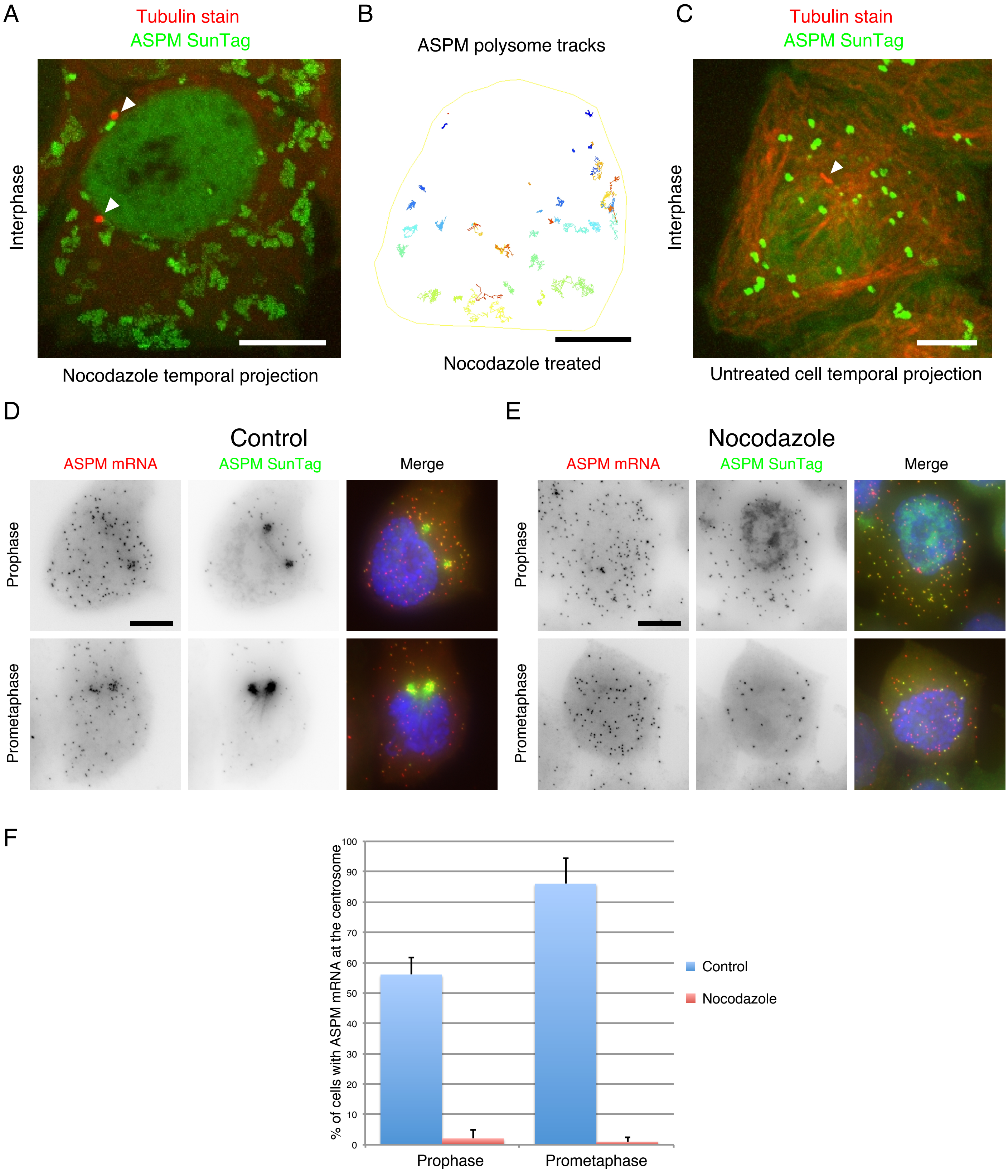


Figure S10

Safieddine et al.

**Figure S10 (related to Figure 7): The effects of nocodazole on ASPM polysome dynamics and ASPM mRNA localization.**

(A) Micrograph represents a temporal projection of SunTagx32-ASPM cells expressing scFv-sfGFP and imaged live during interphase, with labeled MT. The SunTag signal is shown in green and corresponds to ASPM polysomes and mature proteins; the far-red signal is shown in red and corresponds to a tubulin staining. Scale bar: 10 microns. White arrowhead indicates centrosomes.

(B) A TrackMate overlay of the same cell as in A, showing polysomes tracks. Color code represents displacement (dark blue lowest, red highest). The outer yellow outline represents the cell border.

(C) Same legend as in A, but for a control cell untreated with nocodazole.

(D) Micrographs of SunTagx32-ASPM cells expressing scFv-sfGFP and imaged at early mitosis. Left and red: Cy3 fluorescent signals corresponding to tagged and untagged ASPM mRNAs labeled by smFISH with probes against the endogenous mRNA; middle and green: GFP signals corresponding to SunTagx32-ASPM polysomes and mature protein. Blue: DNA stained with DAPI. Scale bar: 10 microns.

(E) Same legend as E, but with cells treated with nocodazole.

(F) Histogram depicting the percentage of cells showing centrosomal localization of ASPM mRNA in early mitosis with and without a nocodazole treatment (n=25 cells per phase, repeated twice). Data represent the mean and standard deviation.

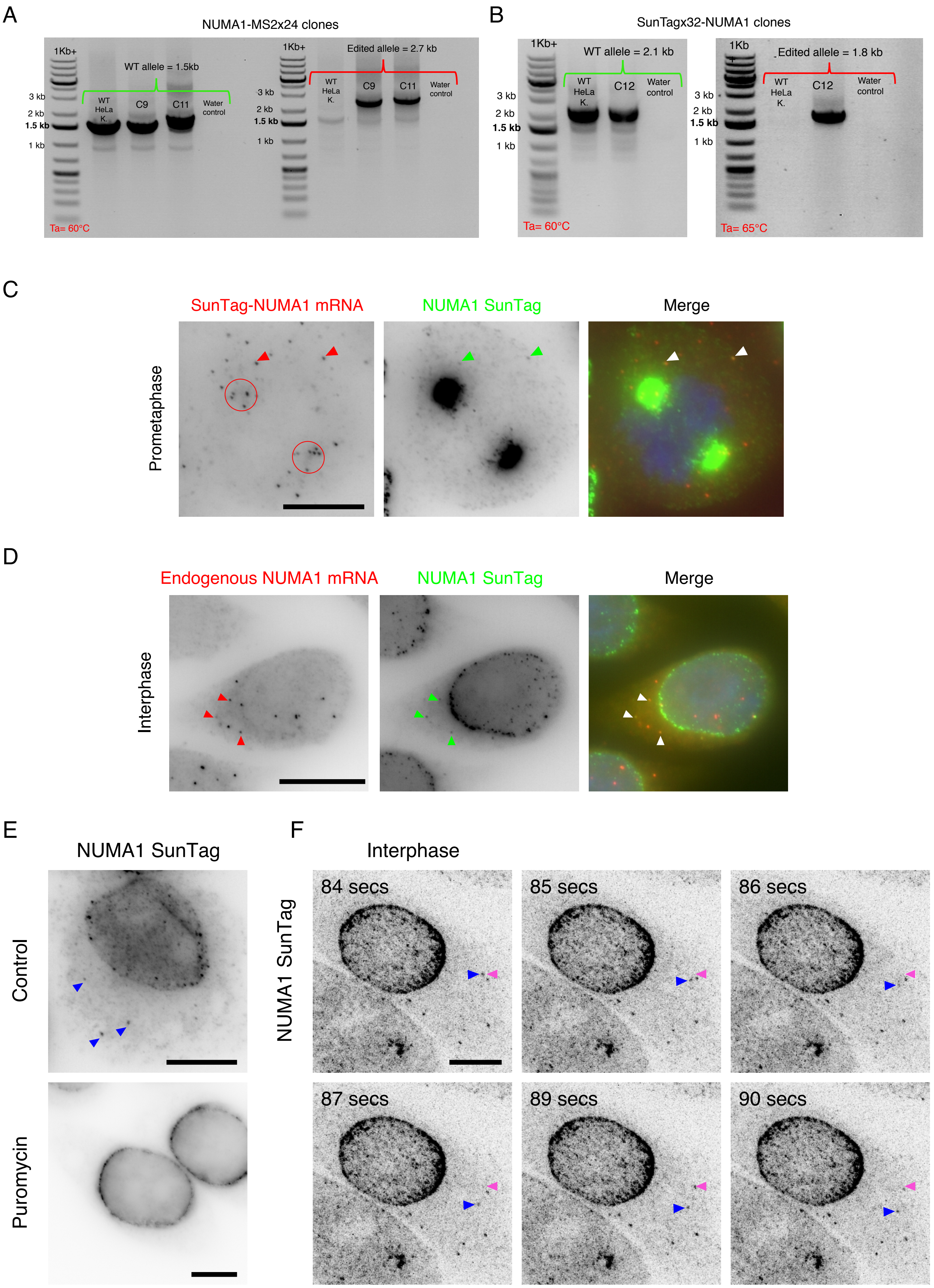


Figure S11

Safieddine et al.

**Figure S11 (related to Figure 8): Characterizing NUMA1-MS2x24 and SunTagx32-NUMA1 clones**

(A) Image is a scan of a gel loaded with the product of a PCR performed on genomic DNA extracted from various NUMA1-MS2x24 clones. Wild-type and edited alleles are differentially amplified and give a product size of 1.5 and 2.7 kbs respectively. A ladder is placed on the left of each amplification with the corresponding size markers. WT HeLa K.: PCR performed on the parental unedited cell line. Water control: PCR performed without any DNA. Ta: annealing temperature.

(B) Image is a scan of a gel loaded with the product of a PCR performed on genomic DNA extracted from a SunTagx32-NUMA1 clone. Wild-type and edited alleles are differentially amplified and give a product size of 2.1 and 1.8 kbs respectively. A ladder is placed on the left of each amplification with the corresponding size markers. WT HeLa K.: PCR performed on the parental unedited cell line. Water control: PCR performed without any DNA. Ta: annealing temperature.

(C) Micrographs of a SunTagx32-NUMA1 cells expressing the scFv-sfGFP. Left and red: Cy3 fluorescent signals corresponding to SunTagx32-NUMA1 mRNAs, revealed by smFISH against the SunTag and puromycin sequences; middle and green: GFP signals corresponding to the SunTagx32-NUMA1 mature protein and polysomes. Blue: DNA stained with DAPI. Scale bar: 10 microns. The red and green arrows indicate an mRNA and a polysome respectively. The red circles indicate clusters of tagged NUMA1 mRNA.

(D) Micrographs of SunTagx32-NUMA1 cells expressing the scFv-sfGFP and images during interphase. Left and red: Cy3 fluorescent signals corresponding to tagged and untagged NUMA1 mRNAs labeled by smiFISH with probes against the endogenous mRNA; middle and green: GFP signals corresponding to SunTagx32-NUMA1 polysomes and mature proteins. Blue: DNA stained with DAPI. Scale bar: 10 microns. Red and green arrowheads indicate SunTagx32-NUMA1 mRNAs and polysomes, respectively. White arrows indicate the overlay of red and green arrows.

(E) Micrographs of SunTagx32-NUMA1 cells, with or without a puromycin treatment and imaged during interphase. The SunTag signal is shown in black. Blue arrowheads indicate polysomes. Scale bars represent 10 microns.

(F) Snapshots of a living SunTagx32-NUMA1 cell imaged during interphase. The SunTag signal is shown in black and corresponds to ASPM polysomes and mature proteins. Scale bar: 10 microns. Time is in seconds. White arrowheads indicate the starting position of an mRNA molecules, while blue one follows its position at the indicated time.

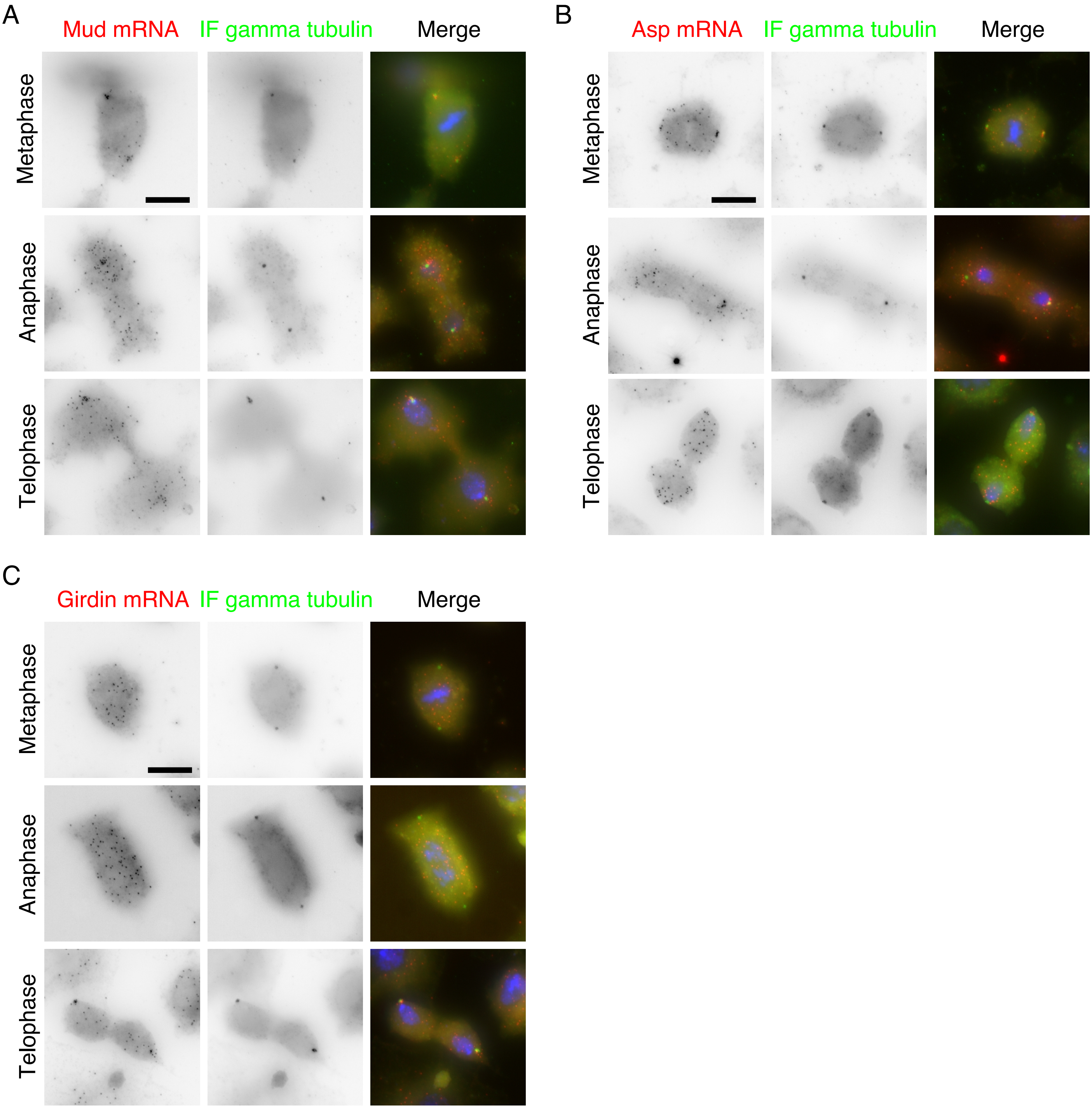
A
Mud mRNA
IF gamma tubulin
Merge
Metaphase
Anaphase
Telophase
B
Asp mRNA
IF gamma tubulin
Merge
Metaphase
Anaphase
Telophase
C
Girdin mRNA
IF gamma tubulin
Merge
Metaphase
Anaphase
Telophase

Figure S12

**Figure S12 (related to Figure 9): Cell cycle dependent centrosomal mRNA localization in S2R+ cells**

(A) Images are micrographs of S2R+ cells during metaphase, anaphase, and telophase. Left and red: Cy3 fluorescent signals corresponding to Mud mRNAs labeled by smiFISH; middle and green: fluorescent signals corresponding to the gamma tubulin protein revealed by IF. Blue: DNA stained with DAPI. Scale bar: 10 microns.

(B) Legend as in A, but for Asp mRNA.

(C) Legend as in A, but for Girdin mRNA.
